# Human lymphatic organoids reconstruct convergent developmental routes to functional lymphatic vessels

**DOI:** 10.64898/2026.11.03.764135

**Authors:** Anmin Wang, Yunpei Zhang, Zilan Li, Karen Wang, Colin P. Johnson, Donghui Xu, Wusiman Maihemuti, Nicole M. Pek, Alonso Tapia Ramos, Kentaro Iwasawa, Varun Sekar, Kavya Pandrangi, Prisha Jain, Takanori Takebe, Chen Yuan Kam, Mingxia Gu

**Affiliations:** Department of Anesthesiology and Perioperative Medicine, David Geffen School of Medicine, Eli and Edythe Broad Center of Regenerative Medicine and Stem Cell Research, University of California, Los Angeles; Los Angeles, CA 90095, USA; Division of Dermatology, Department of Medicine, University of California, Los Angeles; Los Angeles, CA 90095, USA; Broad Stem Cell Research Center, University of California, Los Angeles; Los Angeles, CA 90095, USA; Department of Molecular, Cell and Developmental Biology, University of California, Los Angeles; Los Angeles, CA 90095, USA; Division of Pulmonary Biology, Perinatal Institute, Cincinnati Children’s Hospital Medical Center (CCHMC), Cincinnati, OH, 45229, USA; Center for Stem Cell and Organoid Medicine (CuSTOM), CCHMC, Cincinnati, OH, 45229, USA; Division of Developmental Biology, CCHMC, Cincinnati, OH, 45229, USA; College of Medicine, University of Cincinnati, College of Medicine, Cincinnati, OH, 45267; Division of Gastroenterology, Hepatology and Nutrition, Cincinnati Children’s Hospital Medical Center, Cincinnati, OH 45229, USA; Division of Gastroenterology, Hepatology and Nutrition, Cincinnati Children’s Hospital Medical Center, Cincinnati, OH 45229, USA; Human Biology Research Unit, Institute of Integrated Research, Institute of Science Tokyo (Science Tokyo), Yushima, Bunkyo-ku, Tokyo, Japan; Organoid Medicine Project, T-CiRA Joint Program, Fujisawa, Japan; Center for Stem Cell & Organoid Medicine (CuSTOM), Cincinnati Children’s Hospital Medical Center, Cincinnati, OH 45229, USA; Division of Developmental Biology, Cincinnati Children’s Hospital Medical Center, Cincinnati, OH 45229, USA; Department of Genome Biology, Graduate School of Medicine, The University of Osaka, Suita, Osaka 565-0871, Japan; Premium Research Institute for Human Metaverse Medicine (WPI-PRIMe), The University of Osaka, Suita, Osaka 565-0871, Japan

## Abstract

The lymphatic vasculature maintains fluid balance, regulates immune surveillance, and supports tissue repair, yet how human lymphatic vessels arise and acquire specialized functions remains poorly understood. Here, guided by single-cell and spatial analyses of human embryos, we found that both lateral plate and paraxial mesoderm converge on venous endothelial intermediates with cardinal vein-like characteristics prior to lymphatic specification. Informed by these developmental trajectories, we generated three-dimensional lymphatic organoids from human pluripotent stem cells through two distinct mesodermal routes. The organoids formed capillary-like vascular networks with emerging button-like junctions and distinct endothelial and stromal programs. The lymphatic organoids were transcriptionally benchmarked to human fetal lymphatic endothelium. The organoids supported immune-cell interactions and antigen accumulation and, following transplantation, exhibited interstitial tracer uptake and transport. These findings establish a platform for investigating how distinct developmental origins shape human lymphatic specification, vascular organization, and function, with applications to lymphatic disease modeling and tissue repair.

## Main Text

The lymphatic vasculature is an integral component of nearly every organ, coordinating fluid homeostasis, immunity, and tissue repair. Beyond returning interstitial fluid to the circulation and absorbing dietary lipids, lymphatic vessels regulate immune responses (*1, 2*), support the clearance of macromolecules from cerebrospinal and brain interstitial fluids (*3*), and contribute to heart and bone regeneration (*4, 5*). These diverse functions depend on lymphatic endothelial cells (LECs) acquiring specialized identities and assembling into vessels with distinct structures and transport properties. Disruption of lymphatic function can lead to fluid accumulation and persistent inflammation (*1, 6*), while lymphatic remodeling in cancer can facilitate tumor dissemination (*7*). Nevertheless, therapies that selectively restore or modulate lymphatic function remain limited (*8, 9*). A central unresolved question for both lymphatic biology and therapeutic development is how lymphatic endothelial cell identity, tissue organization, and specialized function is established in humans.

Addressing this question requires experimental systems that connect developmental specification with tissue organization and function. Animal models provide organismal context but may not fully reproduce human developmental programs. Human lymphatic-on-chip and engineered vessel models capture selected aspects of lymphatic structure and function, but their reliance on already specified LECs leaves the earlier events that establish lymphatic identity outside their scope (*10, 11, 12*). Pluripotent stem cell-based approaches provide access to earlier differentiation stages and generate LEC-like populations, yet unbiased transcriptomic benchmarking of their developmental intermediates and differentiated populations against human tissue references remains limited (*13, 14, 15*). The remaining challenge is therefore to comprehensively reconstruct human lymphatic development at a single cell resolution, establish how successive cell states correspond to their *in vivo* counterparts, and determine how different cell populations ultimately give rise to organized lymphatic vasculature with specialized functions.

Organoids offer a framework for addressing this challenge. As three-dimensional, self-organizing systems derived from pluripotent stem cells, they can recapitulate key aspects of developmental progression, multicellular organization, and tissue function in an experimentally accessible human setting (*16, 17, 18, 19*). Applying this framework to the lymphatic vasculature requires an understanding of the differentiation stages and cues that generate human lymphatic tissue. Defining these programs would reveal how lymphatic identity emerges and provide the biological foundation for reconstructing this process in an organoid system. The central challenge is therefore twofold: to delineate human lymphatic development and to reconstruct the progression from lineage specification to organized, functional tissue in an organoid system.

Here, we integrate analyses of human embryonic single-cell and spatial transcriptomic datasets to identify lateral plate- and paraxial mesoderm-associated transcriptional trajectories that converge toward venous and subsequently lymphatic endothelial states. Guided by these findings, we generate lymphatic organoids from human pluripotent stem cells through both mesodermal routes. Time-resolved single-cell profiling reveals cardinal vein-like endothelial populations before lymphatic specification. The resulting organoids acquire a shared lymphatic endothelial identity, supported by single-cell benchmarking against human fetal references, while retaining distinct endothelial transcriptional programs and stromal composition. The organoids form multicellular vascular networks with lymphatic-like junctional architecture, support immune-cell interactions and antigen accumulation. They also enable stage-specific perturbation of developmental signaling to examine how differentiation cues regulate the acquisition of lymphatic identity. Following transplantation into immunocompromised mice, organoid-derived vessels persist and demonstrate drainage function. Together, these findings advance our understanding of human lymphatic development and establish an organoid system for investigating how developmental programs generate lymphatic tissue organization and specialized function. This platform provides a foundation for studying lymphatic dysfunction and the broader roles of lymphatic vessels in immunity, regeneration, and cancer.

## Results

### Spatial and single-cell transcriptomic analyses of early human lymphatic development

Studies in animal models suggest both venous and non-venous origins of LECs (*20, 21, 22*), raising the question of whether human LECs arise exclusively from the cardinal veins or also from other sources. To explore these potential origins, we first examined the spatial distribution of human LEC populations relative to the cardinal veins. We focused on Carnegie stage (CS) 12, when PROX1-positive endothelial cells have been observed in and around the human cardinal veins, preceding lymphatic sprouting at CS13 (*23*). We reanalyzed published human embryonic spatial transcriptomic data (*24*) and annotated candidate cardinal vein regions based on anatomical position and gene expression (Fig. 1A). Because *PROX1* is a key marker of lymphatic endothelial identity but is also expressed in non-endothelial tissues, including the developing liver (*25, 26*), we distinguished *PROX1*⁺ spots with co-detection of *CDH5* or *PECAM1* (CD31) transcripts from *PROX1*⁺ control spots in which none of pan-endothelial markers was detected.

**Fig. 1.**
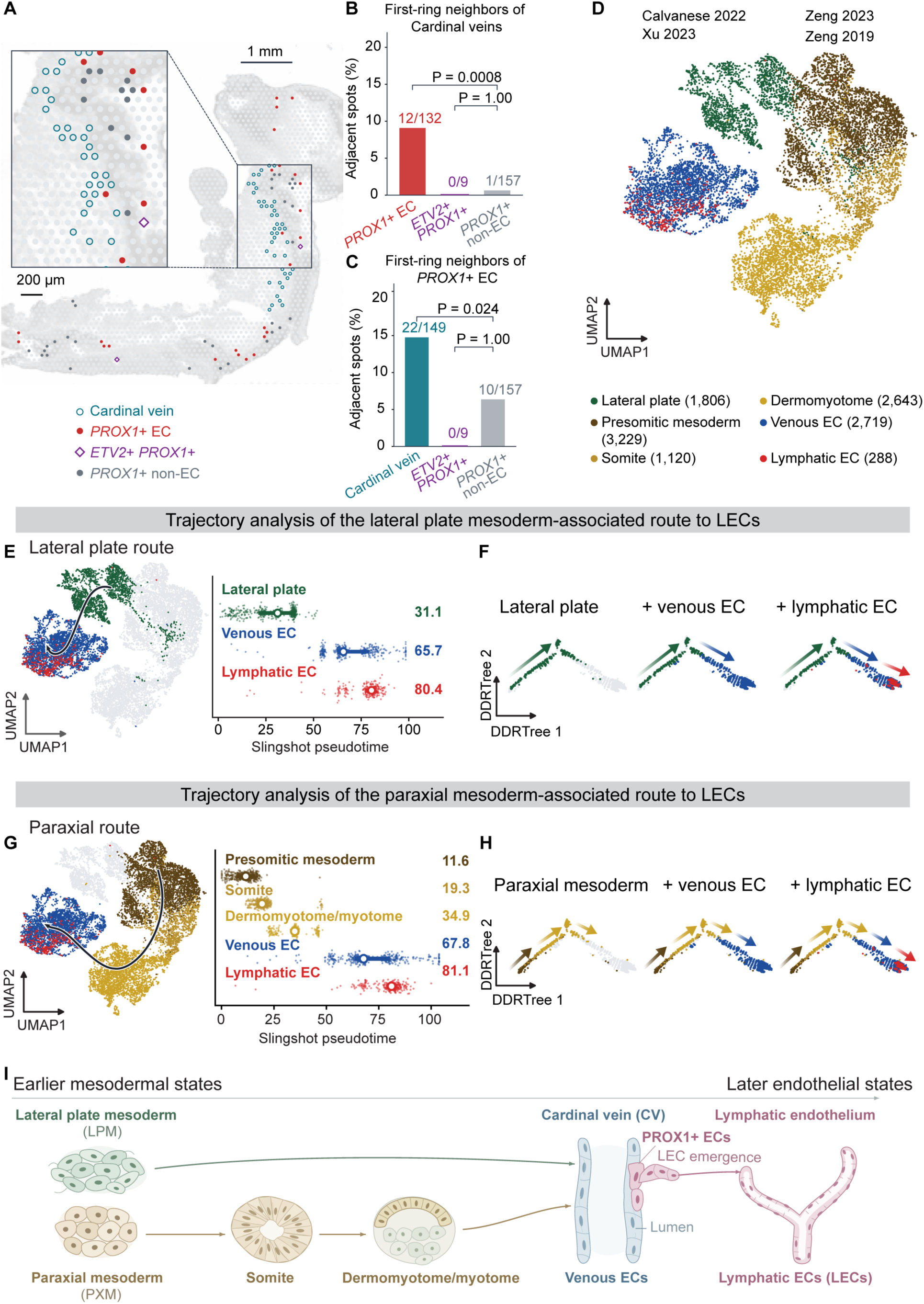
Human embryonic spatial and single-cell analyses identify two mesoderm-associated routes toward lymphatic endothelium. **(A)** Spatial transcriptomic mapping of a Carnegie stage (CS) 12 (CS12) human embryonic trunk section (*24*) (section 1 of eight sections from the analyzed embryo). Candidate cardinal-vein territories were annotated from anatomical position and endothelial/venous gene expression. Open cyan circles: candidate cardinal-vein spots; red circles: *PROX1*⁺ *CDH5*/*PECAM1*⁺ endothelial-associated spots outside the annotated vein; purple diamonds: *ETV2*⁺*PROX1*⁺ spots without detected *CDH5* or *PECAM1*; and gray circles: *PROX1*⁺ spots lacking detectable pan-endothelial markers (regional non-endothelial controls). The inset enlarges the boxed region. Spatial capture spots represent multicellular regions and are not equivalent to individual cells. Scale bars, 1 mm (overview) and 200 µm (inset). (**B**) Fraction of *PROX1*⁺ endothelial-associated spots, *ETV2*⁺*PROX1*⁺ endothelial-marker-negative spots, and *PROX1*⁺ non-endothelial control spots containing at least one candidate cardinal-vein spot out of six directly adjacent spots. Data are pooled across the six spatial sections containing sufficient cardinal vein spots: 12/132 (9.1%), 0/9 (0%), and 1/157 (0.6%), respectively. Two-sided Fisher’s exact tests versus the non-endothelial control: P = 0.0008 and P = 1.00, respectively. (**C**) Reciprocal neighborhood analysis: fraction of candidate cardinal-vein spots, *ETV2*⁺*PROX1*⁺ endothelial-marker-negative spots, and regional control spots with at least one *PROX1*⁺ endothelial-associated spot among their six first-ring neighbors. Across the six sections, fractions were 22/149 (14.8%), 0/9 (0%), and 10/157 (6.4%), respectively. Two-sided Fisher’s exact tests versus the regional control: P = 0.024 and P = 1.00, respectively. (**D**) Integrated single-cell RNA-sequencing analysis of human embryonic mesodermal and endothelial populations at CS7 to CS12 from four studies (*24, 27, 28, 29*). The uniform manifold approximation and projection (UMAP) displays lateral plate mesoderm (n = 1,806), presomitic mesoderm (n = 3,229), somite (n = 1,120), dermomyotome/myotome (n = 2,643), venous endothelial cells (n = 2,719), and lymphatic endothelial cells (n = 288), totaling 11,805 displayed cells. (**E**) Slingshot trajectory analysis of the lateral plate mesoderm-associated route, displayed on the integrated UMAP (left). Right, route-specific pseudotime distributions for lateral plate mesoderm, venous endothelium, and lymphatic endothelium; open circles and adjacent numbers mark median pseudotime, and horizontal bars show the interquartile range. Pseudotime was rescaled from 0 to 100 within the route. (**F**) Monocle 2 DDRTree reconstruction of the lateral plate route using the indicated sequentially added cell populations. Arrows summarize inferred progression along pseudotime. (**G**) Slingshot trajectory analysis of the paraxial mesoderm-associated route, encompassing presomitic mesoderm, somite, dermomyotome/myotome, venous endothelium, and lymphatic endothelium. UMAP trajectories and class-specific pseudotime distributions are displayed as in (E), with pseudotime rescaled within the paraxial route. (**H**) Monocle 2 DDRTree reconstruction of the paraxial route, displayed as in (F). (**I**) Summary schematic of the two candidate developmental routes from lateral plate and paraxial mesoderm toward lymphatic endothelial identity, both involving venous-associated endothelial states. The model summarizes transcriptomic trajectory and spatial-neighborhood associations. Spatial neighborhood comparisons in (B) and (C) used two-sided Fisher’s exact tests with each capture spot as the counted observation; counts were pooled across six sections. P values are reported in the individual panel descriptions.

Spatial neighborhood analysis showed that *PROX1*⁺ endothelial-associated spots were more frequently adjacent to candidate cardinal vein regions than were non-endothelial control spots (12/132, 9.1%, versus 1/157, 0.6%; P = 0.0008; Fig. 1B). Reciprocally, candidate cardinal vein spots were more frequently adjacent to *PROX1*⁺ endothelial-associated spots than were control spots (22/149, 14.8%, versus 10/157, 6.4%; P = 0.024; Fig. 1C). These findings demonstrate a spatial association between emerging LECs and the cardinal veins in early human embryos. Given recent evidence showing paraxial mesoderm-derived *ETV2*⁺*PROX1*⁺ progenitors can directly generate LECs in mice (*22*), we also examined spots co-detecting *ETV2* and *PROX1* but lacking *CDH5* and *PECAM1* (Fig. 1, A to C). Only nine spots exhibited this marker combination, and none were adjacent to *PROX1*⁺ EC spots or cardinal vein regions (Fig. 1, B and C), suggesting that ETV2^+^PROX1^+^ progenitors alone offer a limited representation of emerging LECs.

To identify putative mesodermal progenitors of LECs prior to venous specification, we focused on lateral plate and paraxial mesodermal lineages, based on the spatial association of early LECs with embryonic veins and previous evidence of distinct mesodermal contributions to lymphatic development in mice (*21, 22*). We integrated four published human embryonic single-cell RNA-sequencing datasets spanning CS7 to CS12 (*24, 27, 28, 29*). These datasets were selected from an assembled reference of 41 studies based on developmental stage, representation of LEC-relevant trunk or paraxial tissues, and sequencing methodology. The resulting integrated reference comprised six annotated populations: lateral plate mesoderm (LPM), presomitic mesoderm, somite, dermomyotome/myotome, venous endothelial cells (VECs), and LECs (Fig. 1D and Fig. S1, A to K).

Trajectory inference using Slingshot and Monocle 2 (*30, 31*) revealed distinct transcriptional trajectories connecting mesodermal populations to venous and lymphatic endothelial states. Along the lateral plate-associated route, LPM cells occupied early pseudotime positions, followed by VECs and subsequently LECs (Fig. 1, E and F). Similarly, the paraxial-associated route positioned presomitic mesoderm, somite, and dermomyotome/myotome before venous and lymphatic endothelial populations (Fig. 1, G and H). We further examined the distribution of *ETV2*⁺*PROX1*⁺ cells in the single-cell reference and identified eight double-positive cells among early paraxial populations, including four presomitic mesoderm cells, one somite cell, and three dermomyotome cells. These cells occupied early pseudotime positions without forming a distinct continuous trajectory toward LECs (Fig. S2, A to F). Together, these analyses delineated lateral plate and paraxial mesodermal trajectories toward early human LECs, both involving a venous-associated transcriptional intermediate (Fig. 1I). Building upon mouse studies that identified lineage-biased and region-specific mesodermal contributions (*21, 22, 32*), our findings suggest that both human mesodermal populations could give rise of LECs through a common venous-associated cell state. This proposed model provides a developmental framework for reconstructing lymphatic specification from distinct mesodermal origins *in vitro*.

### Developmentally guided generation of lymphatic organoids from two mesodermal routes

To reconstruct these developmental programs *in vitro* and experimentally validate the computationally inferred developmental trajectories identified in Fig. 1, we established two human pluripotent stem cell (hPSC)-based differentiation strategies to generate paraxial mesoderm-derived lymphatic organoids (PLOs) and lateral plate mesoderm-derived lymphatic organoids (LLOs).

For the paraxial route, cells were sequentially directed through paraxial mesoderm, somite, and dermomyotome-associated stages before endothelial differentiation (Fig. 2A), following established developmental principles (*33, 34*). The hPSC-derived PLOs exhibited robust PAX3 (key paraxial mesoderm marker) staining with minimal HAND1 (key LPM marker) expression, indicating highly efficient and specific paraxial mesoderm specification (Fig. 2B).

**Fig. 2.**
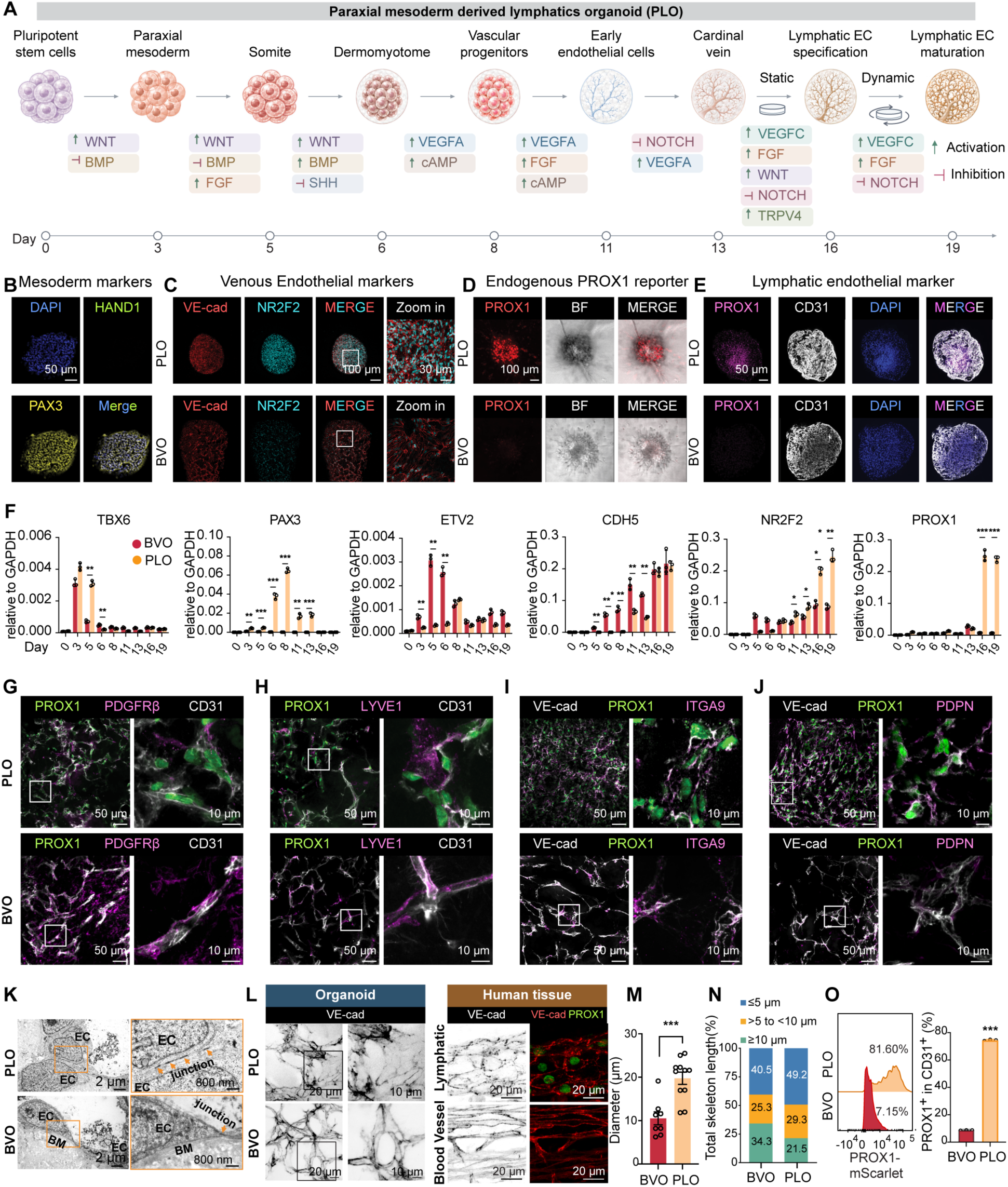
Generation and structural characterization of paraxial mesoderm-derived lymphatic organoids (PLOs). **(A)** Schematic of differentiation of human induced pluripotent stem cells (iPSCs) into PLOs through paraxial mesoderm (day 3), somite (day 5), dermomyotome (day 6), vascular progenitors (day 8), early endothelium (day 11), a cardinal vein-like endothelial stage (day 13), lymphatic endothelial specification (day 16), and maturation (day 19). Stage-specific activation or inhibition of WNT, BMP, FGF, SHH, VEGFA, cAMP, NOTCH, VEGFC, and TRPV4-associated signaling is shown. Arrowheads indicate activation; blunt-ended lines indicate inhibition. **(B)** Representative immunofluorescence staining for HAND1 and PAX3 in day 5 PLOs, with DAPI nuclear counterstaining. Scale bar, 50 µm. **(C)** Representative VE-cadherin (VE-cad) and NR2F2 immunofluorescence in day 13 PLOs and conventional blood vessel organoids (BVOs; non-lymphatic vascular controls). Zoom in panel shows boxed regions with NR2F2^+^VE-cadherin^+^ endothelial structures. Scale bars, 100 µm (overviews) and 30 µm (enlargements). **(D)** Endogenous PROX1-mScarlet reporter fluorescence, corresponding bright-field images, and merged images of PLOs and BVOs one day after initiation of lymphatic induction. Reporter experiments were performed using an H1 human embryonic stem cell line carrying an endogenous PROX1-mScarlet reporter. Scale bar, 100 µm. **(E)** Representative immunofluorescence staining for PROX1 and CD31 in day 19 PLOs and BVOs, with DAPI nuclear counterstaining. Scale bar, 50 µm. **(F)** RT-qPCR time course of *TBX6*, *PAX3*, *ETV2*, *CDH5*, *NR2F2*, and *PROX1* during PLO and BVO differentiation (days 0 to 19). Transcript levels were normalized to *GAPDH*; bar colors distinguish PLOs (orange) and BVOs (red). Data are mean ± SEM. PLO and BVO values were compared using two-sided Student’s t tests. **(G to J)** Representative immunofluorescence comparing PLO and BVO endothelial networks (day 19): **(G)** PROX1, PDGFRβ, and CD31; **(H)** PROX1, LYVE1, and CD31; **(I)** VE-cadherin, PROX1, and ITGA9; and **(J)** VE-cadherin, PROX1, and PDPN. Enlarged views correspond to boxed regions. Scale bars, 50 µm (overviews) and 10 µm (enlargements). **(K)** Transmission electron micrographs of endothelial structures in PLOs and BVOs at day 19. Right panels show enlargements of the boxed regions. EC, endothelial cell; BM, basement membrane. Orange arrows indicate endothelial cell–cell junctions. Scale bars, 2 µm (left) and 800 nm (right). **(L)** Representative VE-cadherin staining in day 19 PLOs and BVOs (left), compared with lymphatic and blood vessels in human embryonic mesenteric tissue as tissue references (right). Enlargements show discontinuous VE-cadherin-positive junctional patterns in PLOs and human lymphatic vessels, in contrast to the more continuous patterns observed in BVOs and human blood vessels. Scale bars, 20 µm (organoid overviews and human tissues) and 10 µm (organoid enlargements). **(M)** Quantification of vessel diameter in BVOs and PLOs from the same differentiation experiment as in (L) (µm). Data are mean ± SEM. Groups were compared using a two-sided Student’s t test. **(N)** Distribution of skeletonized VE-cadherin^+^ junctional segments in BVOs and PLOs from the same differentiation experiment as in (L), categorized by segment length (≤5 µm, >5 to <10 µm, and ≥10 µm) and expressed as percentages of total VE-cadherin^+^ skeleton length. The corresponding percentages were 40.5%, 25.3%, and 34.3% in BVOs and 49.2%, 29.3%, and 21.5% in PLOs, respectively. **(O)** Representative flow-cytometry profiles of endogenous PROX1-mScarlet reporter intensity in PLOs and BVOs (day 19) (left), with quantification of PROX1 reporter-positive cells among CD31⁺ endothelial cells (right). Representative gated fractions were 81.60% in PLOs and 7.15% in BVOs. The gating workflow is shown in Fig. S6A. Reporter experiments used the H1 PROX1-mScarlet line. Data are mean ± SEM. Groups were compared using a two-sided Student’s t test. *P < 0.05, **P < 0.01, ***P < 0.001, ****P < 0.0001; ns, not significant.

To identify signaling cues that could guide the subsequent venous and lymphatic endothelial differentiation, we analyzed signaling-associated transcriptional programs in human embryonic endothelial populations (Fig. S3A). Compared with a common endothelial progenitor reference, venous endothelial cells exhibited reduced VEGF-A, NOTCH, TGF-β, and MEK-ERK signaling signatures, suggesting that inhibition of these pathways may favor venous endothelial specification (Fig. S3B). In contrast, lymphatic endothelial cells showed increased VEGF-C, WNT, and shear/flow-associated signatures, together with reduced NOTCH signaling, suggesting that VEGF-C and WNT activation, NOTCH inhibition, and mechanical cues associated with fluid flow may promote lymphatic differentiation (Fig. S3B). While previous studies have established roles for VEGF-A, NOTCH, TGF-β, and ERK signaling in arterial-venous specification (*35, 36, 37, 38*) and VEGF-C in lymphatic development (*39*), our analysis further identified WNT activation, NOTCH suppression, and shear/flow-associated signaling as potential regulators of lymphatic endothelial differentiation. These findings provided a developmental framework for designing stage-specific differentiation strategies incorporating both biochemical and biomechanical cues.

Guided by these predictions, we performed stage-specific experimental screening to optimize venous and subsequent lymphatic endothelial differentiation. To determine the optimal timing of endothelial induction, we initiated differentiation at successive stages of paraxial patterning. Induction beginning on day 6 yielded the highest CD31 expression among the tested conditions (Fig. S4A, a to c). We next systematically screened combinations of reduced VEGFA exposure and inhibition of NOTCH, TGF-β, and MEK-ERK signaling to promote venous endothelial specification. Reduced VEGFA combined with NOTCH inhibition produced the strongest venous-transcriptional signature, characterized by increased expression of venous markers, including *NRP2*, *APLNR*, *EPHB4*, and *NR2F2* (Fig. S4B, a to b). Notably, additional TGF-β inhibition markedly reduced the venous score, while combined TGF-β and MEK-ERK inhibition also failed to improve venous specification, indicating that selective modulation of VEGFA and NOTCH signaling was sufficient to achieve the most favorable venous endothelial profile. Consistent with these screening results, day 13 lymphatic organoids (corresponding to the cardinal vein stage) exhibited robust NR2F2 expression in VE-cadherin^+^ endothelial networks, whereas NR2F2 was largely absent in blood vessel organoids (BVOs) generated at a comparable differentiation stage using a previously established protocol (*40*). This supported the selective acquisition of venous endothelial identity in our differentiation system (Fig. 2C).

We next optimized lymphatic endothelial induction by screening WNT-associated modulation, continued NOTCH inhibition (RO4929097), PGE2, and TRPV4 activation (GSK1016790A) in the presence of VEGF, guided by our computational analysis (Fig. S3B) and previous studies of lymphatic specification and lymphangiogenesis (*41, 42, 43, 44*). Combined WNT-associated modulation, NOTCH inhibition, and TRPV4 activation yielded the highest lymphatic endothelial marker score (Fig. S4C, a to b). To further investigate the predicted role of shear/flow-associated signaling, we tested mechanical stimulation and pharmacological modulation of PIEZO1- and YAP-associated pathways (*45, 46*). Unexpectedly, static culture produced the strongest PROX1 expression (Fig. S4C.c), suggesting that additional mechanical stimulation was not required to maximize lymphatic marker expression at this stage.

To directly visualize lymphatic endothelial development, we used an H1 human embryonic stem cell line carrying an endogenous PROX1-mScarlet reporter. By day 19, PLOs exhibited robust reporter fluorescence and extensive, interconnected PROX1⁺CD31⁺ endothelial networks, whereas BVOs showed no detectable PROX1 expression (Fig. 2, D and E). Time-course gene-expression analysis further demonstrated sequential induction of paraxial mesodermal markers *TBX6* and *PAX3*, followed by endothelial markers *ETV2* and *CDH5*, and subsequently *PROX1* in PLOs, whereas BVOs lacked both paraxial mesodermal marker induction and subsequent *PROX1* expression (Fig. 2F). The PROX1⁺ endothelial networks also expressed additional lymphatic markers, including LYVE1, ITGA9, and PDPN (Fig. 2, H to J). In contrast to BVOs, which contained abundant PDGFRβ⁺ pericytes, PLOs exhibited limited PDGFRβ staining (Fig. 2G), further highlighting their distinct cellular compositions.

Transmission electron microscopy revealed apposed endothelial membranes forming discrete cell-cell junctions in PLOs, whereas BVOs exhibited more prominent basement membrane and extracellular matrix deposition at endothelial interfaces (Fig. 2K). VE-cadherin staining further revealed discontinuous, button-like junctional features in PLOs, resembling those observed in human lymphatic vessels, in contrast to the more continuous, zipper-like junctions in BVOs and human blood vessels (Fig. 2L). PLOs also exhibited larger vessel diameters than BVOs (Fig. 2M). Quantitative analysis revealed a greater proportion of short VE-cadherin-positive junctional segments in PLOs than in BVOs, consistent with emerging button-like junctions (Fig. 2N). These features were less pronounced than in mature lymphatic vessels, likely reflecting the early developmental stage of PLOs. Flow cytometry further confirmed robust lymphatic specification, with 81.60% of CD31⁺ endothelial cells expressing the PROX1 reporter in representative PLO samples, compared with 7.15% in BVO controls (Fig. 2O).

We next established the lateral plate differentiation route by modifying the conventional BVO protocol with sequential venous and lymphatic induction steps (*40*) (Fig. 3A). Early LLO cultures exhibited robust HAND1 staining with minimal PAX3 expression, indicating efficient and specific lateral plate mesoderm specification (Fig. 3B). Similar to the paraxial route, reduced VEGFA combined with NOTCH inhibition yielded the strongest venous-marker profile (Fig. S5, A and B). Consistent with these screening results, NR2F2 expression was detected within VE-cadherin-positive endothelial networks, confirming the acquisition of venous endothelial characteristics (Fig. 3C). Subsequent lymphatic induction screening again identified combined WNT-associated modulation, NOTCH inhibition (RO4929097), and TRPV4 activation (GSK1016790A) as the condition producing the highest lymphatic endothelial gene signature (Fig. S5, C and D).

**Fig. 3.**
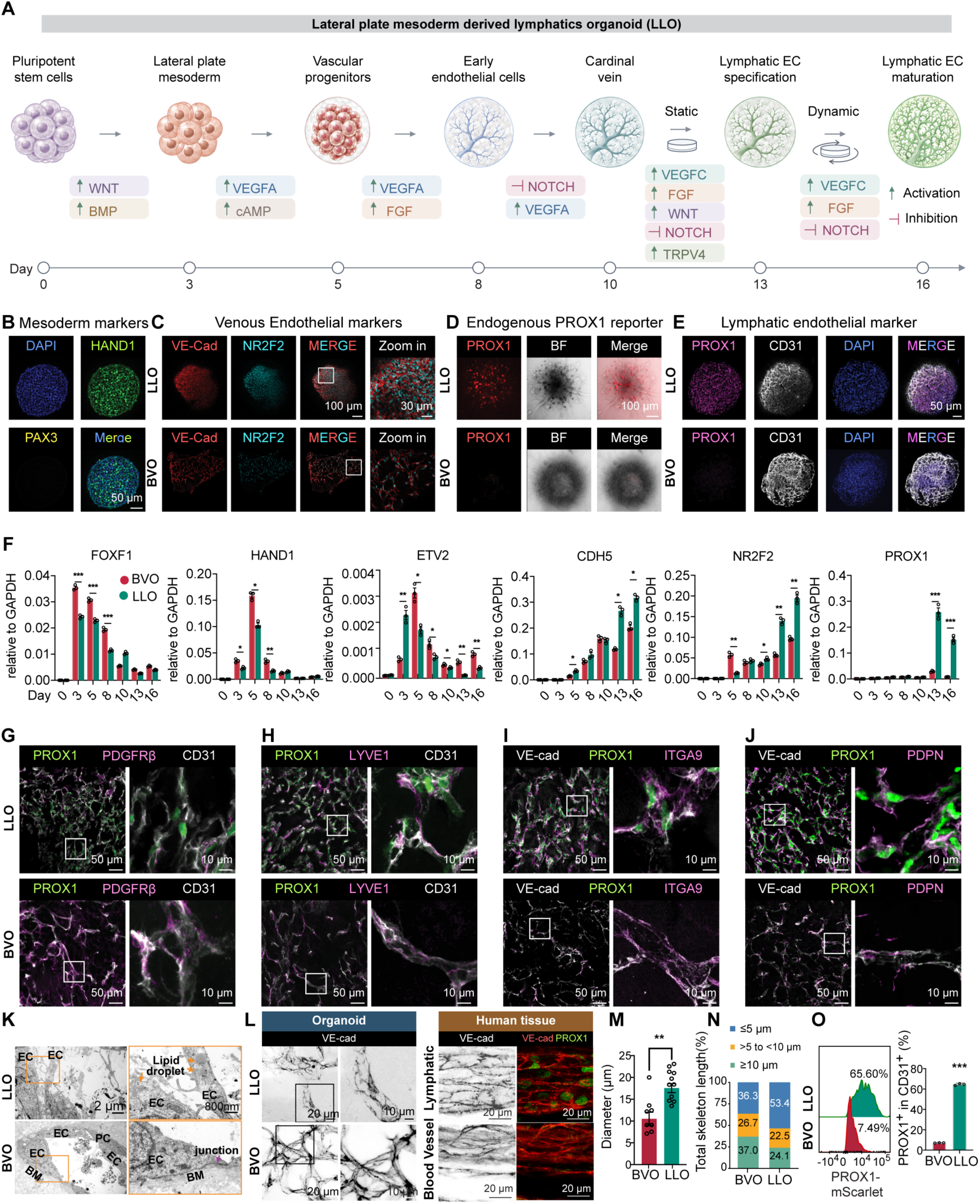
Generation and structural characterization of lateral plate mesoderm-derived lymphatic organoids (LLOs). **(A)** Schematic of stepwise differentiation of human iPSCs into LLOs through lateral plate mesoderm (day 3), vascular progenitors (day 5), early endothelium (day 8), a cardinal vein-like endothelial stage (day 10), lymphatic specification (day 13), and maturation (day 16). Stage-associated activation or inhibition of WNT, BMP, VEGFA, cAMP, FGF, NOTCH, VEGFC, and TRPV4-associated signaling is indicated. Arrowheads indicate activation; blunt-ended lines indicate inhibition. **(B)** Representative immunofluorescence staining for HAND1 and PAX3 in day 3 LLOs, with DAPI nuclear counterstaining. Scale bar, 50 µm. **(C)** Representative VE-cadherin and NR2F2 immunofluorescence in day 10 LLOs and BVO controls. Zoom in panel shows boxed regions with NR2F2^+^VE-cadherin^+^ endothelial structures. Scale bars, 100 µm (overviews) and 30 µm (enlargements). **(D)** Endogenous PROX1-mScarlet reporter fluorescence, corresponding bright-field images, and merged images of LLOs and BVOs one day after initiation of lymphatic endothelial induction. Reporter experiments were performed using an H1 human embryonic stem cell line carrying an endogenous PROX1-mScarlet reporter. Scale bar, 100 µm. **(E)** Representative immunofluorescence staining for PROX1 and CD31 in day 16 LLOs and BVOs, with DAPI nuclear counterstaining. Scale bar, 50 µm. **(F)** RT-qPCR time course of *FOXF1*, *HAND1*, *ETV2*, *CDH5*, *NR2F2*, and *PROX1* during differentiation of LLOs (green) and BVOs (red; days 0 to 16), normalized to *GAPDH*. Data are mean ± SEM. LLO and BVO values were compared using two-sided Student’s t tests. **(G to J)** Representative immunofluorescence comparing LLO and BVO networks for **(G)** PROX1, PDGFRβ, and CD31; **(H)** PROX1, LYVE1, and CD31; **(I)** VE-cadherin, PROX1, and ITGA9; and **(J)** VE-cadherin, PROX1, and PDPN. Enlargements show the boxed regions. Scale bars, 50 µm (overviews) and 10 µm (enlargements). **(K)** Transmission electron micrographs of endothelial structures in LLOs and BVOs at day 16. Right panels show enlargements of the boxed regions. EC, endothelial cell; PC, pericyte; BM, basement membrane. Orange arrows indicate putative lipid droplets; the purple arrow indicates an endothelial cell–cell junction. Scale bars, 2 µm (left) and 800 nm (right). **(L)** Representative VE-cadherin staining in day 16 LLOs and BVOs (left), compared with lymphatic and blood vessels in human embryonic mesenteric tissue as tissue references (right). Enlargements show discontinuous VE-cadherin^+^ junctional patterns in LLOs and human lymphatic vessels, in contrast to the more continuous patterns observed in BVOs and human blood vessels. Scale bars, 20 µm (human tissues); scale bars for organoid images are indicated in the panels. **(M)** Quantification of vessel diameter in BVOs and LLOs from the same differentiation experiment as in (L) (µm). Data are mean ± SEM. Groups were compared using a two-sided Student’s t test. **(N)** Distribution of skeletonized VE-cadherin^+^ junctional segments in BVOs and LLOs from the same differentiation experiment as in (L), categorized by segment length (≤5 µm, >5 to <10 µm, and ≥10 µm) and expressed as percentages of total VE-cadherin-positive skeleton length. The corresponding percentages were 36.3%, 26.7%, and 37.0% in BVOs and 53.4%, 22.5%, and 24.1% in LLOs, respectively. **(O)** Representative flow-cytometry profiles of endogenous PROX1-mScarlet intensity in LLOs and BVOs (left) and the proportion of PROX1 reporter-positive cells among CD31⁺ endothelial cells (right). Representative gated fractions were 65.60% in LLOs and 7.49% in BVOs. The gating strategy appears in Fig. S6B. Reporter experiments used the H1 PROX1-mScarlet line. Data are mean ± SEM. Groups were compared using a two-sided Student’s t test. *P < 0.05, **P < 0.01, ***P < 0.001, ****P < 0.0001; ns, not significant.

By day 16, LLOs exhibited robust PROX1 reporter fluorescence and extensive, interconnected PROX1⁺CD31⁺ endothelial networks, whereas BVOs showed no detectable PROX1 expression by imaging (Fig. 3, D and E). Time-course gene-expression analysis demonstrated early induction of lateral plate mesodermal markers *FOXF1* and *HAND1*, followed by endothelial markers *ETV2* and *CDH5*, venous-associated *NR2F2*, and subsequently *PROX1*. In contrast, BVOs showed lower NR2F2 expression at later stages and minimal PROX1 induction (Fig. 3F). LLOs also expressed additional lymphatic markers, including LYVE1, ITGA9, and PDPN, while exhibiting limited PDGFRβ staining compared with pericyte-rich BVOs (Fig. 3, G to J). Ultrastructural analysis revealed endothelial cell-cell junctions and notable intracellular lipid droplet accumulation in LLOs, whereas BVOs exhibited more prominent basement membrane-associated structures (Fig. 3K). Similar to PLOs, LLOs displayed discontinuous, lymphatic-like VE-cadherin junctions, larger vessel diameters, and a higher proportion of short junctional segments compared with BVOs (Fig. 3, L to N). Flow cytometry further confirmed lymphatic specification, with 65.60% of CD31⁺ endothelial cells expressing the PROX1 reporter in representative LLO samples, compared with 7.49% in BVO controls (Fig. 3O). Representative PLO and LLO gating workflows are shown in Fig. S6A and Fig. S6B, respectively.

Together, these results demonstrate that two distinct mesodermal differentiation strategies generate lymphatic endothelial networks through venous-associated intermediates. Despite their different developmental origins, both PLOs and LLOs acquired robust lymphatic endothelial identity and formed three-dimensional vascular networks exhibiting characteristic lymphatic structural features, including emerging button-like junctions.

### Time-resolved analysis of route-specific venous programs and cardinal vein-like endothelial states

To validate the developmental identities established by the two differentiation strategies and characterize the intermediate cell populations at successive stages, we performed time-resolved single-cell RNA sequencing of BVOs, LLOs, and PLOs prior to lymphatic induction (Fig. 4A). Integration of 110,192 cells identified populations corresponding to LPM, paraxial mesoderm, somite, angioblast, endothelium, and mural/fibro-stromal cells (Fig. 4, B and C). Analysis of cellular composition across differentiation stages revealed distinct developmental patterns (Fig. 4D): BVOs and LLOs were predominantly associated with LPM, whereas PLOs were enriched in paraxial mesoderm and somitic populations (Fig. 4E). These differences were further supported by lineage-associated markers, including *FOXF1*, *HAND1*, *MSX2*, and *PRRX1* in lateral plate populations and *SOX2*, *NKX1-2*, *HES7*, and *WNT3A* in paraxial populations (Fig. 4F). Together, these findings confirmed that the two differentiation protocols established distinct mesodermal programs before endothelial specification.

**Fig. 4.**
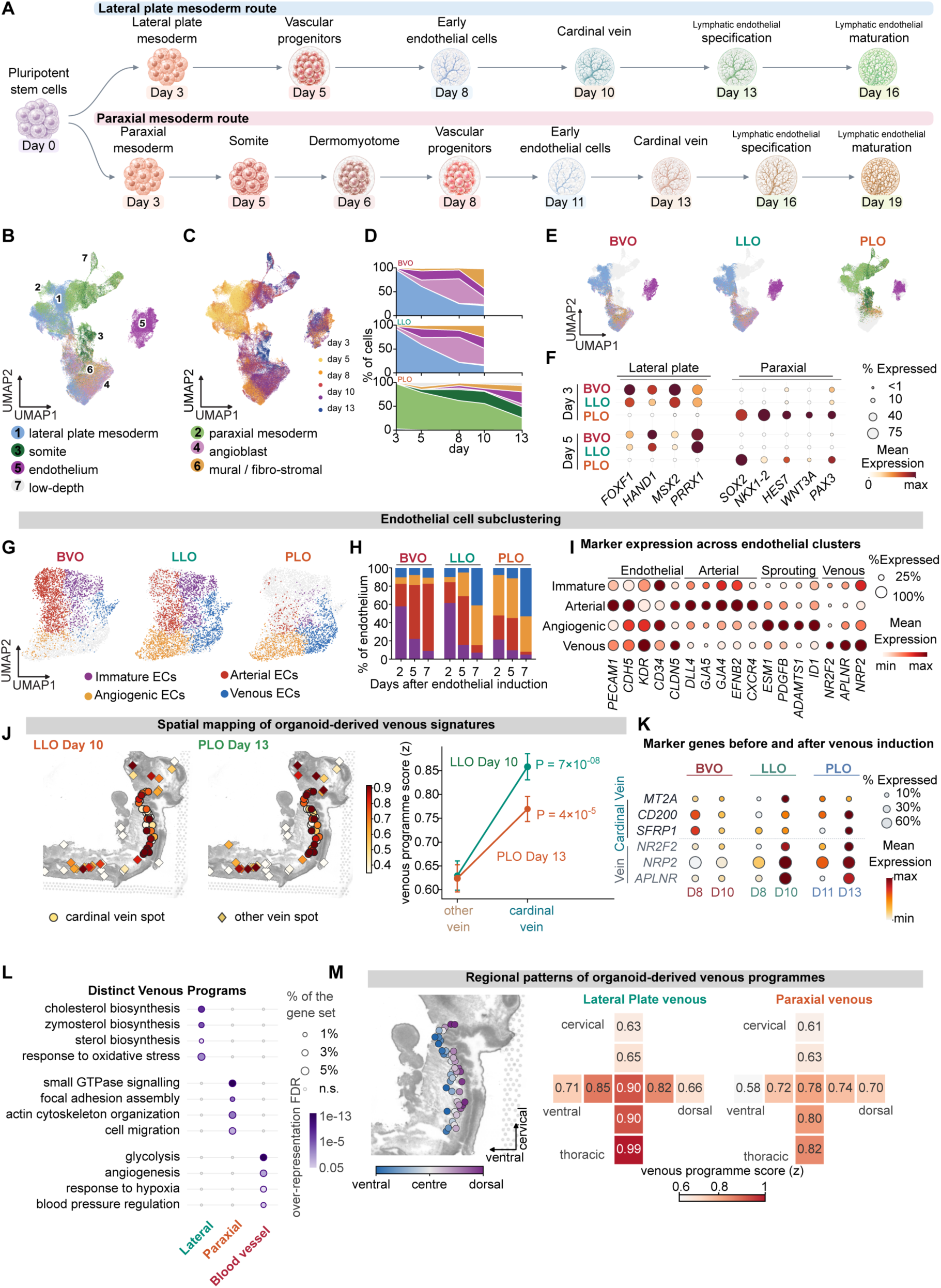
Time-resolved single-cell analysis of the two organoid routes and their cardinal vein-like endothelial programs. **(A)** Differentiation protocol for the lateral plate route (BVOs and LLOs) and paraxial route (PLOs). Days label the indicated protocol stages; sampled days are shown in (C). **(B)** UMAP of an integrated time-series single-cell RNA-sequencing dataset from BVO, LLO, and PLO differentiations, colored by major annotated classes: lateral plate mesoderm, paraxial mesoderm, somite, angioblast, endothelium, mural/fibro-stromal cells, and low-depth/other cells. The comparable developmental window comprises 110,192 cells (BVO and LLO through day 10; PLO through day 13). Numeric labels in the UMAP embedding correspond to the key: 1, lateral plate mesoderm; 2, paraxial mesoderm; 3, somite; 4, angioblast; 5, endothelium; 6, mural/fibro-stromal cells; 7, low-depth cells. **(C)** The same integrated UMAP colored by sampled differentiation day (days 3, 5, 8, 10, and 13, as indicated). **(D)** Relative proportions of the major cell classes at successive sampled days in BVO, LLO, and PLO time courses. **(E)** The integrated UMAP split by organoid type, displaying cells from each organoid in color against the remaining cells in gray. **(F)** Dot plot of lateral plate-associated (*FOXF1*, *HAND1*, *MSX2*, *PRRX1*) and paraxial-associated (*SOX2*, *NKX1-2*, *HES7*, *WNT3A*) genes at early sampled differentiation stages. Dot size represents the percentage of expressing cells; color represents per-gene scaled mean expression. **(G)** UMAP of the endothelial subset, displayed separately for BVOs, LLOs, and PLOs, with cells classified as immature, arterial, angiogenic, or venous endothelial states. **(H)** Endothelial-state composition of BVOs, LLOs, and PLOs at 2, 5, and 7 days after endothelial induction, allowing alignment of route-specific differentiation schedules. **(I)** Dot plot of endothelial identity markers across immature, arterial, angiogenic, and venous endothelial states. Dot area indicates the fraction of cells expressing each gene, and color indicates mean expression scaled within genes. **(J)** Spatial projection of organoid-derived venous transcriptional programs onto a CS12 human embryonic trunk section, comparing cardinal vein-associated spots with spots from other embryonic veins (left). Right, venous-program scores in the two territories for LLO day 10 and PLO day 13. For 126 paired spot comparisons, the mean cardinal vein-other vein score differences were +0.228 for LLO (95% bootstrap confidence interval, 0.16 to 0.30; P = 7 × 10^-8^) and +0.146 for PLO (0.07 to 0.22; P = 4 × 10^-5^). Values are mean ± SEM. Comparisons used two-sided paired Wilcoxon signed-rank tests on 126 matched spot pairs. Confidence intervals were obtained from 5,000 bootstrap resamples. **(K)** Dot plot of candidate cardinal vein-associated markers (*MT2A*, *CD200*, *SFRP1*) and canonical venous genes (*NR2F2*, *NRP2*, *APLNR*) before and after venous induction: BVO and LLO days 8 and 10, and PLO days 11 and 13. Dot area indicates the proportion of expressing cells and color shows per-gene scaled mean expression. **(L)** Gene Ontology biological process over-representation analysis of transcriptional programs distinguishing matched LLO and PLO venous endothelial populations and BVO venous endothelial controls. Dot size indicates gene-set overlap; dot color indicates multiple-testing-adjusted enrichment significance (false-discovery rate). Filled and open circles distinguish significant and nonsignificant terms as shown in the legend. **(M)** Spatial patterning of the lateral plate- and paraxial-associated venous programs within candidate cardinal-vein territories. Left, spatial positions of manually annotated vein spots along the ventral-dorsal axis; right, heatmaps of program scores across anatomical regions and positions, including cervical and thoracic levels. Color indicates standardized venous-program scores across anatomical regions and positions.

One intriguing finding in our study was that both mesodermal differentiation routes converged on venous endothelial identity, raising a key question of whether they generated the same venous endothelial population or distinct venous subtypes reflecting their different developmental origins. To investigate this, we performed endothelial-specific re-clustering, which identified immature, arterial, angiogenic, and venous endothelial states (Fig. 4G). To account for differences in differentiation timing, we compared their relative proportions at matched intervals following endothelial induction (Fig. 4H). BVOs progressively acquired a predominantly arterial endothelial identity, whereas LLOs showed a marked increase in venous endothelial cells during differentiation. PLOs also generated substantial venous endothelium but retained a larger angiogenic endothelial population at the corresponding stage. Marker expression further supported the identities of these endothelial states (Fig. 4I). Thus, although both lymphatic differentiation routes promoted venous endothelial specification, they exhibited distinct endothelial-state compositions during the transition.

We next asked whether the organoid-derived venous populations shared molecular features with the embryonic cardinal vein, a key source of early lymphatic endothelial progenitors (*23*). However, the lack of established cardinal vein-specific markers made it difficult to distinguish this population from other embryonic veins. To address this, we revisited human embryonic datasets to characterize the molecular features of the cardinal vein. By comparing spatially annotated candidate cardinal vein regions with other embryonic venous territories and integrating endothelial single-cell profiles, we identified *MT2A*, *SFRP1*, and *CD200* as candidate cardinal vein-associated markers (Fig. S7, A to H).

Using this molecular reference, we then projected the transcriptional signatures of LLO day-10 and PLO day-13 venous endothelial populations onto the human embryonic spatial data (Fig. 4J). Both populations showed significantly greater correspondence with candidate cardinal vein territories than with other embryonic venous regions, with paired score differences of 0.228 for LLOs (P = 7 × 10^-8^) and 0.146 for PLOs (P = 4 × 10^-5^). We further compared the expression of candidate cardinal vein-associated markers (*MT2A*, *CD200*, and *SFRP1*) and generic venous markers (*NR2F2*, *NRP2*, and *APLNR*) before and after venous induction (Fig. 4K and Fig. S7, I and J). Following venous induction, both LLOs and PLOs exhibited more prominent expression of cardinal vein-associated markers than BVOs, alongside increased expression of conventional venous markers.

Despite their shared venous identity, the two routes retained distinct molecular programs. LLO-derived venous cells were enriched for cholesterol and sterol biosynthesis and oxidative-stress responses, suggesting metabolic specialization, whereas PLO-derived venous cells showed enrichment of small GTPase signaling, focal adhesion, and actin cytoskeleton organization, consistent with enhanced cell remodeling and migration. In contrast, BVO-derived venous cells exhibited glycolytic, hypoxic, and angiogenic programs (Fig. 4L). Spatial analysis revealed broadly similar regional patterns for both LLO- and PLO-derived venous programs, with relatively strong correspondence in thoracic cardinal vein territories and subtle differences in their spatial distributions (Fig. 4M).

Together, these analyses demonstrate that both mesodermal routes converge on endothelial populations with cardinal vein-like features while retaining distinct cellular compositions and transcriptional programs. These findings provide a foundation for investigating how mesodermal origin influences subsequent lymphatic endothelial specification.

### Distinct endothelial and stromal programs in lymphatic organoids

To further characterize the cellular composition of LLOs and PLOs and assess their transcriptional similarity to human lymphatic vessels, we performed single-cell RNA sequencing of both organoid systems at their differentiation endpoints, alongside conventional BVOs for comparison (Fig. 5). The three organoid systems exhibited distinct cellular compositions (Fig. 5, A to C). BVOs consisted primarily of mesenchymal cells (46%), pericytes (26%), and arterial endothelial cells (20%). In contrast, LLOs were highly enriched in lymphatic endothelial cells (69%), with a smaller mesenchymal population (27%), whereas PLOs contained both lymphatic endothelial (32%) and mesenchymal (68%) populations. Notably, pericytes were abundant in BVOs but largely absent from both lymphatic organoid systems, consistent with the lack of pericyte coverage characteristic of native initial lymphatic capillaries (*47*).

**Fig. 5.**
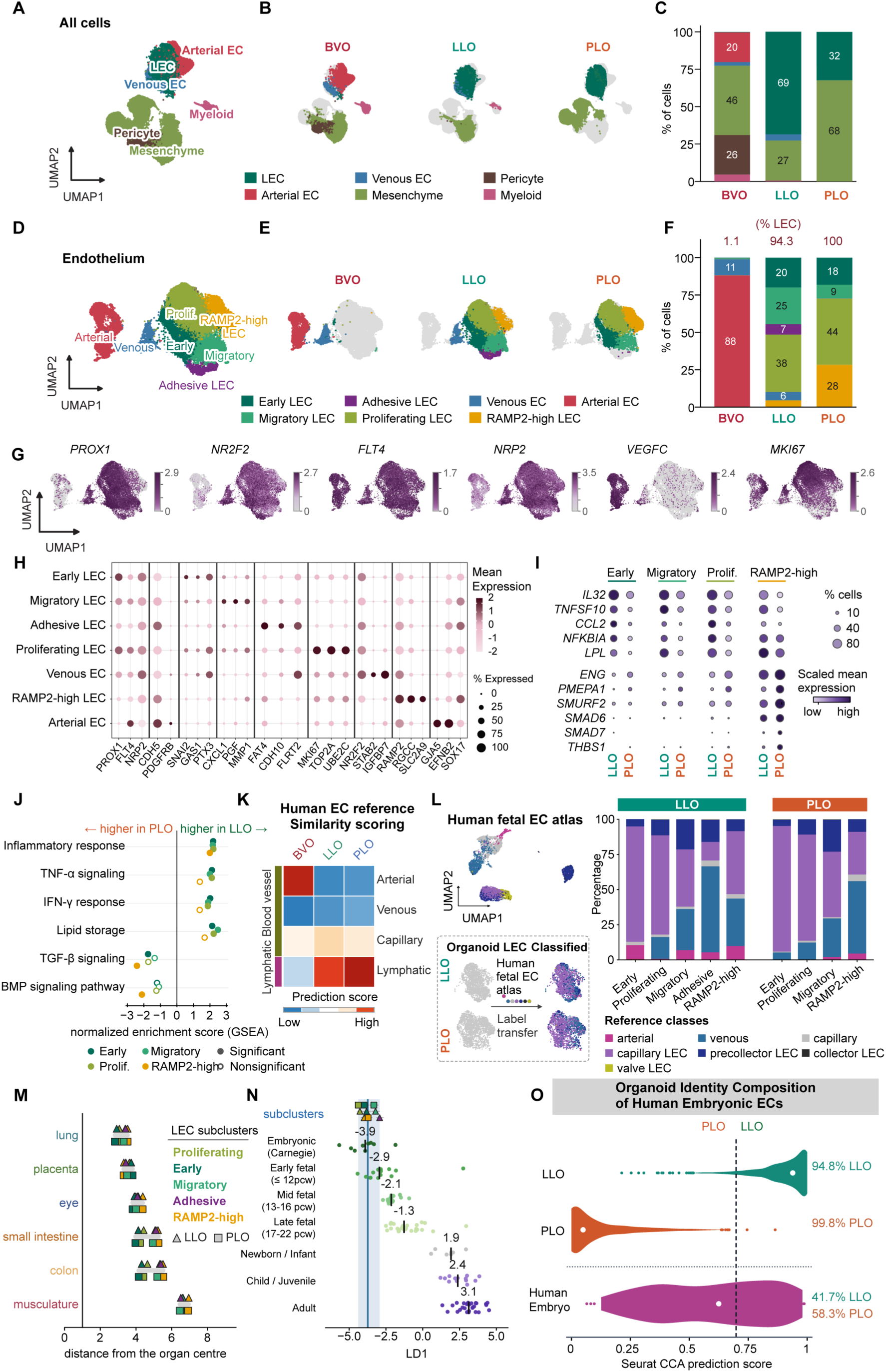
Lymphatic endothelial heterogeneity, route-specific programs, and correspondence to human fetal endothelium. **(A)** Integrated UMAP of 46,284 cells from BVOs, LLOs, and PLOs, showing lymphatic, arterial, and venous endothelial cells together with mesenchymal, pericyte, and myeloid populations. **(B)** The same embedding split by organoid type. Cells belonging to the displayed organoid are colored by class and the other two organoid populations are shown in gray. **(C)** Proportions of the major annotated cell classes in each organoid. Lymphatic endothelial cells constitute approximately 69% of all profiled LLO cells and 32% of PLO cells, whereas the PLO sample contains a larger mesenchymal compartment; the percentages refer to all profiled cells. **(D)** Reclustering of 18,864 endothelial cells from the three organoid types, identifying early, migratory, adhesive, proliferating, and RAMP2-high lymphatic endothelial states, together with arterial and venous endothelial cells. **(E)** The endothelial UMAP split by organoid type, with other cells shown in gray. **(F)** Composition of endothelial subtypes within BVOs, LLOs, and PLOs. The percentage of the endothelial compartment assigned to lymphatic states is shown above each bar: 1.1% in BVOs, 94.3% in LLOs, and 100% in PLOs. These endothelial-normalized percentages are distinct from the proportions of all cells in (C). **(G)** Feature plots of *PROX1*, *NR2F2*, *FLT4*, *NRP2*, *VEGFC*, and *MKI67* expression across endothelial states. Color intensity represents gene expression, scaled separately for each gene. **(H)** Dot plot of lineage and endothelial state-associated genes across seven endothelial states. Dot size shows the percentage of cells detecting each gene; color shows scaled mean expression within each gene. **(I)** Route-associated differential expression of selected genes within corresponding LLO and PLO lymphatic endothelial states (early, migratory, proliferating, and *RAMP2*-high). Genes shown include *IL32*, *TNFSF10*, *CCL2*, *NFKBIA*, *LPL*, *ENG*, *PMEPA1*, *SMURF2*, *SMAD6*, *SMAD7*, and *THBS1*. Dot size indicates the percentage of expressing cells and color indicates scaled mean expression. **(J)** Gene set enrichment analysis of matched LLO and PLO lymphatic endothelial states for inflammatory response, TNF-α/NF-κB, interferon-γ response, lipid storage, TGF-β signaling, and BMP signaling programs. Positive normalized enrichment scores indicate enrichment in LLOs, and negative scores indicate enrichment in PLOs; filled versus open points denote the significance status indicated in the key. **(K)** Similarity scores of BVO, LLO, and PLO endothelial populations against a human endothelial reference encompassing arterial, venous, blood capillary, and lymphatic categories. Color denotes relative prediction/similarity score within the displayed heatmap. **(L)** Reference-based label transfer from a human fetal endothelial atlas comprising 8,530 endothelial cells in 36 donor blocks to organoid-derived lymphatic endothelial states. The schematic illustrates the labeling workflow, and stacked bars show predicted reference-class proportions (including capillary, pre-collecting, collecting, valve, and venous/blood endothelial categories) for early, proliferating, migratory, adhesive, and RAMP2-high states in LLOs and PLOs. Predicted labels indicate transcriptomic similarity to the reference cell types. **(M)** Distances of LLO and PLO lymphatic endothelial subclusters from fetal organ reference centers in a discriminant space, displayed for lung, placenta, eye, small intestine, colon, and musculature. Triangles denote LLO and squares denote PLO subclusters; colors identify endothelial state. Distances are scaled to the reference distribution of each organ. **(N)** Projection of organoid lymphatic endothelial subclusters onto a reference developmental-stage discriminant axis (LD1), including embryonic, fetal, newborn/infant, juvenile, and adult endothelial reference groups. Locations correspond to relative positions along the reference axis rather than an exact gestational age for organoid cells. **(O)** Seurat canonical-correlation-based classification distinguishing LLO and PLO lymphatic endothelial cells. Violin plots show the LLO-class prediction scores for held-out LLO cells, held-out PLO cells, and human embryonic lymphatic endothelial cells. The indicated classifications were 94.8% for LLO, 99.8% for PLO, and 41.7% of embryo LECs assigned to the LLO class. The vertical dashed line indicates the displayed classification threshold. Statistical analysis. Panel (J) reports gene set enrichment statistics using the multiple-testing-adjusted false-discovery rate (significance threshold FDR < 0.05). Panels (A to I) and (K to O) are primarily descriptive or computational comparisons. Percentages in (C) use all cells, whereas those in (F) use endothelial cells only.

To further characterize endothelial identity and heterogeneity, we re-clustered the endothelial compartment and identified seven transcriptionally distinct states, including early, migratory, adhesive, proliferating, and *RAMP2*-high LECs, together with venous and arterial endothelial cells (Fig. 5, D and E). RAMP2 is a component of the adrenomedullin receptor complex. Disruption of this signaling axis impaired lymphatic development in mice (*48*). Notably, LECs accounted for 94.3% of endothelial cells in LLOs and 100% in PLOs, compared with only 1.1% in BVOs (Fig. 5F). The lymphatic populations expressed established LEC markers, including *PROX1*, *FLT4*, and *NRP2* (Fig. 5G). *NR2F2* was enriched in venous endothelial cells but reduced in most LEC populations, with a distinct *NR2F2*-high venous population persisting in LLOs but not detected in PLOs (Fig. 5E,G,H). Additional transcriptional markers distinguished early, migratory, adhesive, proliferating, and *RAMP2*-high LEC states (Fig. 5H).

Despite acquiring a shared LEC identity, LLOs and PLOs retained distinct transcriptional programs. LLO-derived LECs exhibited higher expression of immune- and lipid-associated genes, including *IL32*, *TNFSF10*, *CCL2*, *NFKBIA*, and *LPL*, whereas PLO-derived LECs showed increased expression of *ENG*, *PMEPA1*, *SMAD6*, *SMAD7*, and *THBS1*, genes involved in TGF-β/BMP signaling regulation and extracellular matrix remodeling (Fig. 5I). Consistently, gene-set enrichment analysis revealed stronger inflammatory response, TNF-NF-κB signaling, interferon-γ response, and lipid-storage programs in LLO-derived LECs. In contrast, PLO-derived LECs exhibited greater enrichment of TGF-β- and BMP-associated signaling programs (Fig. 5J). Notably, the enrichment of lipid-associated programs in LLOs was consistent with the intracellular lipid droplets observed by electron microscopy, which were not evident in PLOs (Figs. 2K and 3K). Together, these findings suggest that LLO-derived LECs exhibit greater immune-associated and lipid-handling features, whereas PLO-derived LECs retain stronger developmental signaling and vascular remodeling programs.

To benchmark the identity, tissue specificity, and developmental maturity of organoid-derived LECs against native human lymphatic endothelium, we compared their transcriptional profiles with human endothelial references spanning different organs and developmental stages. Both LLOs and PLOs showed greater similarity to human LECs than BVOs, which more closely resembled arterial endothelial cells (Fig. 5K). Label transfer using human fetal endothelial references further identified predominantly capillary LEC identities in both organoid systems, with no detectable collecting or valve LEC populations at this stage (Fig. 5L). Across tissue references, LLOs and PLOs exhibited broadly similar patterns, with the closest transcriptional correspondence to lung and placental LECs rather than a single organ-specific identity (Fig. 5M). Developmental-stage mapping further placed both organoid systems closest to early embryonic (Carnegie-stage) LECs, rather than later fetal or postnatal populations (Fig. 5N). Finally, to investigate whether the distinct mesodermal route-associated signatures were also present in human embryos, we applied a transcriptional classifier trained to distinguish LLO- and PLO-derived LECs. Human embryonic LECs exhibited a mixture of LLO-like and PLO-like signatures (Fig. 5O), suggesting that the molecular differences associated with the two differentiation routes may also be represented during human lymphatic development.

To investigate whether these endothelial differences were accompanied by distinct supporting microenvironments, we analyzed the non-endothelial populations of each organoid system (Fig. S8, A to D). The LLO non-endothelial compartment consisted predominantly of fibroblasts, whereas PLOs contained substantial stromal and paraxial mesenchymal populations. In contrast, BVOs retained a prominent pericyte population (Fig. S8, E and F). These differences suggested that the two mesodermal differentiation strategies generated distinct multicellular environments surrounding the lymphatic endothelial networks.

We next investigated whether the distinct mesodermal origins of LLOs and PLOs also established different cellular microenvironments supporting lymphatic development. CellChat analysis (*49*) revealed distinct patterns of predicted cell-cell communication, with LLO fibroblasts contributing *ANGPT1*-*TEK* and *SLIT2*-*ROBO1* interactions, whereas PLO stromal and mesenchymal populations exhibited *VEGFC*-, *MDK*-, and *PTN*-associated signaling (Fig. S8, G to J). Strikingly, PLO stromal cells accounted for 100% of the predicted *VEGFC* ligand contribution among the analyzed supporting-cell populations (Fig. S8K). Given the essential role of VEGFC in lymphatic development (*39*), this finding suggests that PLOs may establish an endogenous VEGFC-producing stromal niche that supports lymphatic endothelial development. Together, these findings suggest that distinct mesodermal origins not only shape LEC identities but also establish different supporting-cell environments, potentially influencing how lymphatic development is regulated within each organoid system.

### Immune-cell recruitment, antigen accumulation, and transplantation-associated lymphatic functions

Initial lymphatic capillaries guide CCR7-expressing immune cells, including dendritic cells and T lymphocytes, toward lymphatic vessels through CCL21-dependent signaling (*50, 51*). To determine whether our lymphatic organoids support similar immune-cell interactions, we incubated BVOs, LLOs, and PLOs with human peripheral blood mononuclear cells (PBMCs) and examined the penetration of CCR7-positive cells (Fig. 6A). Both lymphatic organoid systems exhibited significantly increased CCR7-associated fluorescence compared with BVO controls (Fig. 6B), supporting enhanced recruitment and accumulation of CCR7-positive immune cells within the lymphatic organoid environment.

**Fig. 6.**
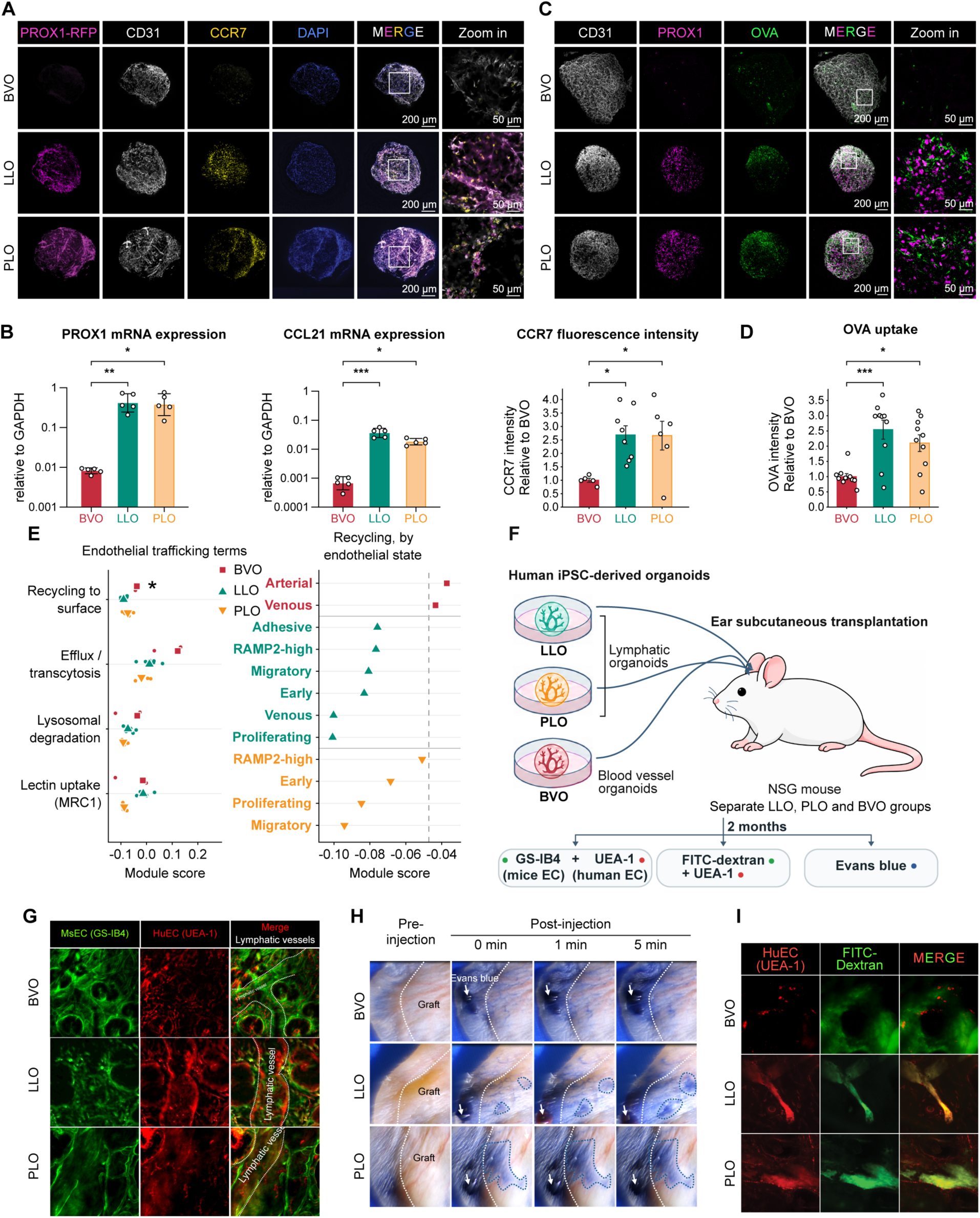
*In vitro* and *in vivo* functional characterization of human lymphatic organoids. **(A)** Representative images of BVOs, LLOs, and PLOs after incubation with human peripheral blood mononuclear cells (PBMCs), stained for endogenous PROX1 reporter fluorescence (PROX1-RFP), CD31, CCR7, and DAPI. Merged views and enlargements of boxed regions show organoid-associated CCR7⁺ cells. Scale bars, 200 µm (overviews) and 50 µm (enlargements). **(B)** RT-qPCR quantification of PROX1 and CCL21 mRNA expression in BVOs, LLOs, and PLOs, normalized to GAPDH (left and middle), and quantification of organoid-associated CCR7 fluorescence intensity relative to BVOs (right). Data are mean ± SEM. Groups were compared using one-way ANOVA with Tukey’s multiple-comparisons test; symbols denote Tukey-adjusted P values. **(C)** Representative fluorescent ovalbumin (OVA) accumulation assay in BVOs, LLOs, and PLOs, showing CD31, PROX1, fluorescent OVA, overlays, and enlargements. Scale bars, 200 µm (overviews) and 50 µm (enlargements). Organoids were incubated with fluorescent OVA for 12 h, washed, and cultured for an additional 3 days before measurement. **(D)** Quantification of organoid-associated fluorescent OVA intensity in BVOs, LLOs, and PLOs. Each dot represents one organoid. Data are mean ± SEM. Groups were compared using one-way ANOVA with Tukey’s multiple-comparisons test; symbols denote Tukey-adjusted P values. **(E)** Transcriptomic endothelial trafficking module scores related to recycling to the plasma membrane, efflux/transcytosis, lysosomal degradation, and MRC1-associated lectin uptake (left). Right, recycling-to-surface module scores for arterial, venous, and the indicated lymphatic endothelial states, including RAMP2-high LECs, grouped by organoid route. The recycling score comparison used a two-sided Mann-Whitney U test comparing BVO (n = 2 states) with LLO/PLO (n = 10 states), without multiple-testing correction. **(F)** Experimental schematic of subcutaneous transplantation of human LLOs, PLOs, and BVOs into the ears of NOD scid gamma (NSG) mice. Grafts were examined approximately two months later using dual lectin labeling to visualize human-associated (UEA-I) and mouse-associated (GS-IB4) vascular structures, local Evans blue tracer imaging, and tracing with 2,000-kDa FITC-dextran. **(G)** Representative lectin labeling of transplanted ear tissues from BVO, LLO, and PLO groups, using GS-IB4 (mouse-associated endothelial structures, green) and UEA-I (human-associated endothelial structures, red). Merged views show the spatial relationship between host and graft-associated vascular structures. Dashed contours indicate the vessel-like regions labeled in the panel. GS-IB4 (20 µg/mL, 5 µL) and ATTO 594-conjugated UEA-I (2 µg/mL, 5 µL) were administered by local subcutaneous injection near the graft two months after transplantation. **(H)** Mouse ears before and at 0, 1, and 5 min after interstitial Evans blue injection. White dotted contours identify graft regions; blue dotted contours delineate the tracer-associated regions shown after injection. White arrows indicate the injection region. Evans blue (5 mg/mL in PBS, 5 µL per imaging site) was administered by local subcutaneous injection. **(I)** Representative images of transplanted tissue labeled with UEA-I (human-associated endothelial structures, red) and interstitially injected FITC-dextran (2,000 kDa; green), with merged images for BVO, LLO, and PLO graft groups. The images show tracer localization relative to graft-associated human endothelial structures. FITC-dextran was administered at 20 µg/mL in a volume of 5 µL by local subcutaneous injection together with ATTO 594-conjugated UEA-I. Transplantation images in (G to I) represent three independent experiments with one mouse per group per experiment (three mice per group). *P < 0.05, **P < 0.01, ***P < 0.001, ****P < 0.0001; ns, not significant.

Beyond immune-cell trafficking, LECs can capture and retain antigens, contributing to antigen archiving in draining lymph nodes (*2, 52*). To assess whether our organoids acquired similar antigen-handling properties, we examined the accumulation of fluorescent ovalbumin (OVA) following prolonged incubation. Both LLOs and PLOs exhibited significantly greater OVA-associated fluorescence than BVOs, particularly in PROX1⁺ endothelial regions (Fig. 6, C and D). To explore potential mechanisms underlying this enhanced antigen accumulation, we analyzed endothelial intracellular trafficking programs. Notably, LLO- and PLO-derived LECs exhibited reduced recycling-to-surface transcriptional signatures compared with BVO endothelial populations (Fig. 6E), suggesting that decreased antigen recycling may contribute to intracellular antigen retention.

To determine whether organoid-derived lymphatic vessels could undergo further maturation and acquire interstitial drainage capacity *in vivo*, we transplanted BVOs, LLOs, and PLOs into the ear pinnae of immunodeficient mice and allowed two months for vascular maturation (Fig. 6F). Species-specific lectin staining revealed connections between UEA-1⁺ human lymphatic networks and GS-IB4⁺ host mouse vessels in LLO and PLO grafts, whereas BVO grafts exhibited limited evidence of vascular integration (Fig. 6G). To assess lymphatic drainage, we injected Evans blue into the ear interstitium, where tracer uptake occurs primarily through lymphatic rather than blood vessels. Serial imaging revealed rapid tracer distribution along branching vessel-like structures within LLO and PLO grafts. In contrast, Evans blue remained largely confined to the injection site in BVO-transplanted ears, with limited tracer spread appearing to occur through the surrounding host lymphatic vessels (Fig. 6H). To further distinguish lymphatic uptake from blood vascular transport, we injected high-molecular-weight (2,000-kDa) FITC-dextran into the ear interstitium, which preferentially enters lymphatic vessels due to its restricted permeability across blood capillaries. FITC-dextran was detected within UEA-1⁺ human endothelial structures in LLO and PLO grafts, but remained largely outside these structures in BVO grafts, providing additional evidence of interstitial tracer uptake by transplanted LLOs (Fig. 6I). Together, these findings suggest that LLO- and PLO-derived vascular networks can persist, mature, and acquire lymphatic-like drainage functions following transplantation, extending their functional properties beyond those observed *in vitro*.

## Discussion

This study establishes a developmentally guided human lymphatic organoid platform that reconstructs successive stages of lymphatic development, from mesodermal specification through cardinal vein-like endothelium to multicellular lymphatic networks with emerging biological functions. By directing human pluripotent stem cells through lateral plate and paraxial mesoderm, we show that both starting populations support lymphatic differentiation while generating distinct endothelial and stromal programs. These findings support a model in which a shared lymphatic identity emerges from different mesodermal programs without erasing route-associated molecular diversity. Unlike models assembled from already specified LECs, this platform provides experimental access to the developmental transitions that establish lymphatic identity within a three-dimensional human tissue environment.

Our findings provide critical insights into the developmental origins of human lymphatic endothelium. Mouse lineage-tracing studies have revealed strong lineage biases and anatomically distinct contributions to lymphatic endothelium, including predominant paraxial contributions across multiple tissues and lateral plate and second-heart-field contributions in specific territories (*21, 32, 53*). This literature encompasses both specification through venous endothelium and direct differentiation from mesoderm-derived progenitors (*21, 22*). Our findings extend these lineage maps by demonstrating that both human lateral plate and paraxial mesoderm can support a common venous-associated program of lymphatic differentiation. Human embryonic analyses identified candidate trajectories from both populations, and time-resolved organoid profiling recovered cardinal vein-like endothelial states before lymphatic specification. This venous-associated progression is consistent with the cardinal vein-associated emergence of LECs in human embryos (*23*), while extending the developmental framework to earlier mesodermal populations. Together, these results highlight the potential diversity of human lymphatic origins while establishing a system for experimentally investigating venous-associated lymphatic specification.

Importantly, despite converging on a shared lymphatic identity, the two differentiation routes retained distinct molecular characteristics. These differences emerged at the venous stage, with LLO-derived cells enriched for metabolic and oxidative-stress-associated programs, whereas PLO-derived cells showed stronger migration and cytoskeletal remodeling signatures. Following lymphatic specification, LLO-derived LECs exhibited greater inflammatory and lipid-associated programs, while PLO-derived LECs were enriched for TGF-β/BMP-associated signaling. Notably, both LLO-like and PLO-like transcriptional signatures were also detected in human embryonic LECs, suggesting that similar molecular diversity exists during human lymphatic development. Together, these findings raise the possibility that distinct mesodermal origins contribute to lymphatic endothelial specialization beyond the acquisition of a shared LEC identity. However, because the two differentiation routes also establish different stromal environments and signaling conditions, future lineage-tracing studies and reciprocal endothelial-stromal cocultures will be important to determine whether these differences reflect developmental origin, local microenvironmental cues, or both.

The two differentiation routes also generated distinct stromal environments, providing an opportunity to study how supporting cells contribute to lymphatic development. Notably, PLO stromal cells expressed VEGFC and showed predicted VEGFC-FLT4/KDR interactions with LECs, suggesting a potential endogenous source of lymphangiogenic support (*39*). In contrast, LLO fibroblasts exhibited predicted ANGPT1-TEK and SLIT2-ROBO1 signaling. These findings suggest that mesodermal origin may shape not only endothelial identity but also the surrounding cellular microenvironment, potentially influencing lymphatic network formation and maintenance. Future studies using selective stromal perturbations under controlled exogenous growth-factor conditions will help establish the functional contributions of these distinct supporting-cell populations.

Beyond transcriptional identity, our findings suggest that lymphatic organoids recapitulate key structural and functional features of developing lymphatic vessels. Discontinuous VE-cadherin junctions and limited pericyte representation resemble features of initial lymphatic capillaries (*47*). Importantly, the ability of these lymphatic organoids to interact with immune cells and accumulate antigens suggests that aspects of lymphatic immune surveillance can emerge before full vascular maturation. Reduced recycling-associated transcriptional programs further suggest a potential mechanism for antigen retention, although long-term antigen archiving, as observed in lymph nodes, remains to be established (*2, 52*). Following transplantation, the acquisition of interstitial tracer uptake suggests further functional maturation *in vivo*. However, their capacity to restore physiological lymphatic drainage and integrate with host lymphatic networks requires further evaluation in mouse models of lymphedema. Another important direction will be to investigate whether LLOs and PLOs can model lymphatic malformations and how their distinct developmental origins may influence disease manifestations.

Several developmental questions also remain. These organoids primarily capture early lymphatic development and lack mature collecting vessels and valves, which may be promoted through prolonged culture and stage-specific mechanical stimulation. Although our study demonstrates that both lateral plate and paraxial mesoderm can generate lymphatic endothelium through venous-associated intermediates *in vitro*, whether venous specification is obligatory and the extent to which each route contributes to lymphatic development *in vivo* remain to be determined. Combining lineage tracing with selective depletion of lymphatic populations derived from each mesodermal route during mouse embryonic development could clarify their respective contributions and reveal potential compensatory mechanisms.

Together, our study establishes a human stem cell-based platform for reconstructing lymphatic development from distinct mesodermal origins to vascular networks with emerging functional properties, providing new opportunities to investigate human lymphatic biology, model lymphatic disorders, and explore the roles of lymphatic vessels in tissue homeostasis and repair.

## Acknowledgments

We thank Dr. Hongkui Deng, Dr. Hasina Outtz Reed, Dr. Esak Lee, Dr. Sathish Srinivasan, Dr. James M. Wells, Dr. Yue Liu, Dr. Jing Wang, Dr. Enbo Zhu and Meiyan Wang for their valuable support and assistance with this study. We acknowledge the UCLA Translational Pathology Core Laboratory for tissue procurement.

## Author contributions

Conceptualization: AW and MG

Methodology: AW

Investigation: AW, YZ, ZL, KW, CPJ, DX, WM, VS, KP, and PJ

Formal analysis: AW

Validation: ZL

Data curation: AW

Software: AW

Resources: KI and TT

Writing – original draft: AW and MG

Writing – review and editing: AW, MG, CPJ, NMP, and ATR

Supervision: MG and CYK

Project administration: AW and MG

Funding acquisition: MG

## Data and materials availability

The sequencing data generated in this study have been deposited in the NCBI Gene Expression Omnibus (GEO) under accession number GSE352484. Additional data supporting the findings of this study are available in the main text and Supplementary Materials. Research materials are available from the corresponding author upon reasonable request.

## Code availability

Analysis scripts and source tables for Figure reproduction are available at https://github.com/AnmWang/Lymphatics. Panel-specific reproduction details and limitations are documented in the repository.

## Supplementary Materials

### Materials and Methods

#### Human embryonic transcriptomic analysis

##### Dataset assembly and developmental stage harmonization

Cells were obtained from a previously assembled human embryonic single-cell transcriptomic reference comprising 41 published datasets. Because developmental ages were reported using different conventions across studies, all available annotations were harmonized to Carnegie stages (CS). Directly reported Carnegie stages were retained, whereas post-conceptional weeks were converted using a day-based developmental reference table. Because developmental ages were frequently rounded to integer weeks in the original studies, conversions resulting in stage intervals were retained as distinct strata rather than assigned to individual stages. Two datasets with ambiguous age annotations were classified as unresolved and excluded.

Studies were selected using four sequential criteria: (1) developmental age convertible to Carnegie stage (35 of 41 datasets); (2) inclusion of cells from CS12 or earlier (8 datasets); (3) availability of trunk or paraxial tissue (6 datasets); and (4) use of droplet-based single-cell sequencing (4 datasets). Relevant dissections were subsequently selected at the sample level (Fig. S1A).

Eleven captures were retained, including the aorta and vitelline artery dissections from Calvanese et al. (*28*), head-upperTrunk and viscera-lowerTrunk captures from Xu et al. (*29*), body and caudal-half captures from Zeng et al. (*27*), and five whole-body captures from Zeng et al. (*24*). Six non-trunk dissections from Calvanese et al. were excluded. Because the retained samples represented whole-embryo or trunk-level dissections rather than organ-specific tissues, no additional visceral tissue exclusions were applied. Sample-level filtering removed 16,661 cells, leaving 48,094 cells for further analysis.

##### Single-cell RNA sequencing quality control

Quality control was performed independently for each sample using thresholds for detected genes, total transcript counts, and mitochondrial transcript proportions. The mitochondrial threshold was defined as five median absolute deviations above the sample-specific median. Hemoglobin-expressing cells co-expressing *GYPA* or *PTPRC* were excluded. Scrublet (*54*) was applied separately to each sample, resulting in the removal of 69 predicted doublets.

Cells were subsequently restricted to mesodermal and endothelial populations. Clusters lacking sufficient marker support for their original annotations were excluded, together with cardiac-associated clusters. The resulting dataset comprised 23,625 cells (Fig. S1, F to I).

For additional assessment of potential doublets, Scrublet and scDblFinder (*55*) were independently applied to the retained cells within each sample. In total, 1,840 cells were flagged by at least one method, and 441 were identified by both methods (Fig. S1J). These additional predictions were retained as quality-control annotations and were not used for further cell exclusion.

Cell-cycle phases were assigned using S-phase and G2/M-associated gene scores, without regression of cell-cycle-associated expression from the data (Fig. S1K). One contributing study contained no mitochondrial genes in its deposited expression matrix; therefore, mitochondrial fractions for those cells were structurally zero and were not considered comparable with the remaining datasets.

##### Data integration and dimensionality reduction

Gene expression matrices were normalized by library size and log-transformed. To reduce study-associated batch effects, Harmony (*56*) was applied using study identity as the batch correction variable, with 50 dimensions retained for downstream analysis.

Uniform manifold approximation and projection (UMAP) was performed using the Harmony-corrected representation, with n_neighbors = 10 and spread = 3. These parameters were used to control the visualization layout and were not interpreted as modifying distances in the corrected representation. A single frozen UMAP embedding was used consistently across the relevant Figures (Fig. 1, D, E, and G; Fig. S1, B to D; Fig. S2A).

##### Cell population annotation and marker expression

Six mutually exclusive cell classes were defined for downstream visualization and analysis: lateral plate mesoderm (1,806 cells), presomitic mesoderm (3,229 cells), somite (1,120 cells), dermomyotome/myotome (2,643 cells), venous endothelial cells (2,719 cells), and lymphatic endothelial cells (288 cells).

Endothelial cells were identified using a pan-endothelial marker gate requiring expression of at least four genes among *CDH5*, *PECAM1*, *EGFL7*, *CLDN5*, *RAMP2*, *ESAM*, *ICAM2*, *SOX18*, *TIE1*, and *ECSCR*, each detected at two or more counts. Within this endothelial compartment, cells expressing *PROX1* at two or more counts were designated lymphatic endothelial cells, whereas the remaining endothelial cells were classified as venous endothelial cells for display purposes. Thus, the venous and lymphatic classes represented complementary subdivisions of a single annotated endothelial compartment rather than populations independently defined by distinct positive marker sets.

The somite class represented the fine-annotation presomitic-to-somite transition population, comprising 1,085 cells reassigned from the presomitic mesoderm class and 35 cells from the dermomyotome class. Three cells expressing *PROX1* above the defined threshold did not satisfy the pan-endothelial gate and were excluded from the endothelial classes. Overall, the six displayed classes contained 11,805 of the 11,808 cells.

Marker expression was summarized using the mean log-normalized expression within each cell class, followed by gene-wise scaling across the six classes. Dot size represented the percentage of cells expressing each marker (Fig. S1E).

##### Spatial transcriptomic analysis

Spatial transcriptomic data were obtained from a CS12 to 13 human embryo, comprising 14,284 capture spots across eight tissue sections. Each capture spot measured 55 µm in diameter and was positioned on a hexagonal lattice with a center-to-center spacing of 100 µm.

Putative lymphatic precursor spots were identified by detectable *PROX1* expression together with *CDH5* or *PECAM1* expression, provided that the spots were located outside the candidate cardinal vein region.

An additional population of *ETV2*⁺*PROX1*⁺CD31⁻*CDH5*⁻ spots was defined by detectable *ETV2* and *PROX1* expression and the absence of detectable *PECAM1* and *CDH5*. This definition was based on the late-angioblast signature described by Lupu et al. (*22*), with *CDH5* included as an additional exclusion marker.

Control spots were defined as *PROX1*-positive spots located outside the candidate cardinal vein region that lacked detectable expression of all ten pan-endothelial markers: *CDH5*, *PECAM1*, *CLDN5*, *EGFL7*, *RAMP2*, *ESAM*, *KDR*, *TIE1*, *SOX18*, and *ERG*. Spots meeting the endothelial double-positive definition were excluded from the control group.

Spatial adjacency was defined using the six directly neighboring capture spots in the first ring of the hexagonal lattice, corresponding to a center-to-center distance of 100 µm. Adjacency analyses were restricted to the six sections containing at least three candidate cardinal vein spots. Group-level adjacency fractions were compared against the control population using two-sided Fisher’s exact tests (Fig. 1, A to C).

Because individual spatial capture spots may contain transcripts from multiple cells, co-detection of endothelial and lymphatic markers within a spot was interpreted as spatial co-localization rather than definitive evidence of co-expression within a single cell.

##### Trajectory inference using Monocle 2

Developmental trajectories were reconstructed using Monocle 2 (*31*) with the DDRTree algorithm applied to the uncorrected expression representation, with ncenter = 50.

The resulting trajectory comprised three major states: an endothelial arm containing 3,299 cells (pseudotime 10.03 to 21.24), a mesodermal root containing 5,928 cells (pseudotime 0.00 to 9.34), and a dermomyotome arm containing 2,581 cells (pseudotime 9.96 to 12.10).

The root state was assigned based on the highest proportion of presomitic mesoderm cells.

For visualization of the proposed developmental routes, individual cell classes were sequentially highlighted on the same fitted trajectory, while cells not yet included in each displayed route were shown in light grey (Fig. 1, F and H; Fig. S2, C to F).

Monocle 2 pseudotime was derived from an uncorrected expression representation containing study-associated shifts and was interpreted as relative developmental ordering.

##### Independent trajectory analysis using Slingshot

Slingshot (*30*) was applied independently to the paraxial and lateral plate routes to evaluate the consistency of inferred developmental ordering.

For each route, 2,000 highly variable genes were selected, followed by principal component analysis using 30 components. The starting population was specified as presomitic mesoderm for the paraxial route and lateral plate mesoderm for the lateral plate route. Terminal populations were not specified, allowing Slingshot to infer one lineage for each route.

Pseudotime was independently rescaled to a range of 0 to 100 for each route. Accordingly, pseudotime values were compared within individual routes and not directly between the paraxial and lateral plate trajectories.

For visualization, fitted principal curves were projected onto the frozen UMAP embedding by smoothing the embedding coordinates against pseudotime. The displayed curves were shortened at both ends, so their visual endpoints did not represent the original fitted endpoints (Fig. 1, E and G; Fig. S2, A and B).

Agreement between Slingshot and Monocle 2 was assessed by comparing developmental ordering. Trajectory analyses do not establish lineage relationships.

##### Identification and trajectory positioning of *ETV2*⁺*PROX1*⁺ cells

*ETV2*⁺*PROX1*⁺ cells were identified using a permissive expression threshold requiring at least one count for each gene.

Eight double-positive cells were identified within the paraxial mesodermal populations, including four presomitic mesoderm cells, one somite cell, and three dermomyotome cells. No cells satisfying these criteria were detected within the lateral plate route.

The position of these eight cells along the inferred trajectory was compared with 2,000 randomly generated sets of eight paraxial cells matched to the same cell-class composition. Pseudotime distributions were additionally compared against all paraxial cells using a two-sided Mann-Whitney test (Fig. 1, G and H; Fig. S2, B and F).

#### Human pluripotent stem cell culture and vascular organoid generation

##### Human pluripotent stem cell maintenance

Human pluripotent stem cell (hPSC) lines, including H1 PROX1: mScarlet (NIH-0043), CDH5-EGFP, H9-GFP (NIHhESC-0062), and BA049, were obtained from Cincinnati Children’s Hospital Medical Center (CCHMC), including the Takebe and Jason laboratories. The BU1-ST (SFTPC-tdTomato) cell line was obtained from Boston University (BU). Cells were maintained under feeder-free conditions in StemFlex medium (Thermo Fisher Scientific, Cat. No. A3349401) at 37°C in a humidified atmosphere containing 5% CO₂. The culture medium was refreshed according to the manufacturer’s instructions, and cells were routinely passaged to maintain their undifferentiated state.

Cells were maintained on six-well plates coated with growth factor-reduced Matrigel diluted 1:50 in DMEM/F12.

##### Embryoid body formation

For the generation of 3D embryoid bodies (EBs), hPSCs were dissociated into single cells using 0.5 mM EDTA and passed through a 40-µm Flowmi cell strainer (SP Bel-Art, Cat. No. H13680-0040). EBs containing approximately 500 to 1,000 cells each were generated using AggreWell 400 plates (STEMCELL Technologies) in aggregation medium supplemented with 20 µM Y-27632 (ROCK inhibitor), according to the manufacturer’s instructions. Following 24 h of incubation, the resulting EBs were harvested, designated as day 0, and washed once with N2B27 basal medium. The EBs were subsequently subjected to either conventional blood vascular organoid or lymphatic vascular organoid differentiation protocols, as described in the following sections.

Wide-bore pipette tips were used during EB handling and medium transfer to minimize mechanical disruption.

##### Medium preparation

EB aggregation medium consisted of 40 mL of KnockOut DMEM/F12, 10 mL of KnockOut Serum Replacement (KOSR), 0.5 mL of GlutaMAX, 0.5 mL of non-essential amino acids (NEAA), 35 µL of β-mercaptoethanol (prediluted 1:100 in PBS), and 0.5 mL of Antibiotic-Antimycotic.

N2B27 basal medium consisted of 250 mL of DMEM/F12, 250 mL of Neurobasal medium, 10 mL of B27 supplement, 5 mL of N2 supplement, 2.5 mL of GlutaMAX, 350 µL of β-mercaptoethanol (prediluted 1:100 in PBS), and 5 mL of Antibiotic-Antimycotic.

Blood vessel induction medium. Blood vessel induction (BVI) medium consisted of StemPro-34 medium supplemented with 15% fetal bovine serum (FBS), 100 ng/mL VEGF-A, and 100 ng/mL FGF2.

##### Extracellular matrix preparation and organoid embedding

A collagen I solution was prepared by combining 750 µL of 0.1 N NaOH, 313 µL of 10× DMEM, 63 µL HEPES, 49 µL of 7.5% NaHCO₃, 31 µL GlutaMAX, 460 µL Ham’s F-12 medium, and 3.33 mL PureCol collagen I. PureCol collagen I was added last, and the solution was gently mixed to minimize bubble formation.

The collagen I solution was subsequently mixed with Matrigel at a 3:1 volume ratio to produce the final embedding matrix.

At the designated embedding stage, EBs were collected and the culture medium was removed before resuspension in the embedding matrix. Individual EBs were transferred using 200-µL wide-bore pipette tips into separate wells of an organoid embedding sheet (STEMCELL Technologies), with each EB embedded in a 20-µL matrix droplet.

The embedding sheet was incubated at 37 °C for 45 min to allow matrix polymerization. The polymerized droplets containing individual organoids were then gently released by washing with culture medium and transferred into ultra-low attachment six-well plates. Each well contained 18 embedded organoids in 3 mL of the corresponding differentiation medium.

##### Generation of conventional blood vessel organoids

Conventional BVOs were generated following the previously described protocol of Wimmer et al. (*40*), beginning with EB formation in AggreWell 400 plates.

##### Generation of lateral plate mesoderm-derived lymphatic organoids (LLOs)

LLOs were generated using a stepwise differentiation protocol designed to promote endothelial specification followed by lymphatic-lineage development.

On day 0, EBs were cultured in N2B27 basal medium supplemented with CHIR99021 and BMP4.

On day 3, the medium was replaced with N2B27 basal medium containing forskolin and VEGF-A.

On day 5, individual spheroids were embedded in collagen I-Matrigel matrix as described above and transferred to ultra-low attachment six-well plates.

On day 8, the medium was replaced with StemPro-34 supplemented with FBS, VEGF-A, FGF2, and RO4929097.

On day 10, the culture medium was replaced with StemPro-34 containing FBS, VEGF-C, FGF2, RO4929097, WNT5B, and GSK1016790A.

On day 13, WNT5B and GSK1016790A were withdrawn, while VEGF-C, FGF2, and RO4929097 were maintained at their previous concentrations in StemPro-34 medium supplemented with FBS. The medium was subsequently refreshed every three days.

##### Generation of PLOs

Paraxial mesoderm-derived lymphatic organoids (PLOs) were generated using a modified differentiation protocol with an initial stage of mesodermal specification followed by endothelial induction and lymphatic-lineage differentiation.

On day 0, EBs were cultured in N2B27 basal medium supplemented with KOSR, LDN193189, CHIR99021, and Y-27632.

On day 1, the culture medium was replaced with the same formulation without Y-27632.

On day 3, FGF2 was added.

On day 4, the medium was replaced with N2B27 basal medium containing KOSR.

On day 5, the medium was replaced with N2B27 basal medium containing KOSR, CHIR99021, GDC-0449, and BMP4.

On day 6, individual spheroids were embedded in collagen I-Matrigel matrix and transferred to ultra-low attachment six-well plates, with 18 embedded spheroids in 3 mL medium per well. The medium consisted of StemPro-34 supplemented with FBS, VEGF-A, and forskolin.

On day 8, the same medium formulation was supplemented with FGF2.

On day 11, the medium was replaced with StemPro-34 containing FBS, VEGF-A, FGF2, and RO4929097.

On day 13, the medium was replaced with StemPro-34 containing FBS, VEGF-C, FGF2, RO4929097, WNT5B, and GSK1016790A.

On day 16, WNT5B and GSK1016790A were withdrawn, whereas VEGF-C, FGF2, and RO4929097 were maintained at their previous concentrations in StemPro-34 medium containing FBS. The medium was subsequently refreshed every three days.

#### Organoid dissociation and single-cell preparation

##### Dissociation of non-embedded vascular organoids

Non-embedded vascular organoids were washed with calcium- and magnesium-free Dulbecco’s phosphate-buffered saline (DPBS) and incubated with TrypLE at 37 °C for 4 min.

Organoids were gently triturated using a P1000 pipette and examined under a microscope. If dissociation was incomplete, incubation was continued with trituration and microscopic assessment at 2-min intervals until a single-cell suspension was obtained. Complete dissociation typically required 7 to 8 min.

##### Dissociation of matrix-embedded vascular organoids

Matrix-embedded vascular organoids were pooled and washed with calcium- and magnesium-free Hank’s balanced salt solution (HBSS).

The digestion solution consisted of 0.5 mg/mL Liberase TH and 5 mg/mL Dispase prepared in RPMI (+/+). The solution was passed through a 0.2-µm filter and maintained on ice before use.

Organoids were incubated in the digestion solution at 37 °C for 30 to 40 min on a rocker operating at 30 rpm. After the initial 5 min, DNase I was added to a final concentration of 60 U/mL.

Organoids were gently triturated every 5 min using a wide-bore P1000 pipette tip. After 15 to 20 min, trituration was continued using a standard P1000 tip. Dissociation was monitored microscopically, and digestion was extended when necessary, provided that cell viability was maintained. Low-retention pipette tips were used when appropriate to minimize cell loss.

Once a single-cell suspension was obtained, enzymatic digestion was terminated by adding cold complete medium containing at least 10% FBS. Cells were collected by centrifugation at 300 × g for 5 min and resuspended in MACS buffer containing 0.5% MACS BSA Stock Solution and 2 mM EDTA in DPBS, or the appropriate buffer for the downstream sequencing workflow.

For sequencing experiments, cells were washed at least twice to remove residual matrix and debris. When necessary, suspensions were passed through a 40-µm cell strainer to remove remaining aggregates. Filtration was omitted when no aggregates, debris, or matrix fragments were visible, to minimize cell loss.

##### Dissociation of transplanted vascular organoids

Transplanted organoids were washed with calcium- and magnesium-free DPBS to remove residual blood. Excised graft-containing tissues were cut into approximately 2 to 3-mm fragments and incubated in 400 IU/mL collagenase IV prepared in calcium- and magnesium-free DPBS. The enzyme solution was passed through a 0.2-µm filter before use.

Digestion was performed at 37 °C for 1 h on a rocker operating at 30 rpm. DNase I was added after the initial 5 min to a final concentration of 60 U/mL. Tissue fragments were gently triturated every 10 min using a wide-bore P1000 tip.

Digestion was terminated using cold complete medium containing at least 10% FBS, followed by centrifugation at 300 × g for 5 min. The resulting pellet was resuspended in RPMI and passed through a 40-µm cell strainer to remove undigested tissue fragments.

Cells in the filtrate were collected by centrifugation at 300 × g for 5 min. When visible erythrocyte contamination remained, the pellet was resuspended in ACK red blood cell lysis buffer and incubated on ice for 5 min, with gentle inversion midway through the incubation. Cells were subsequently collected by centrifugation at 300 × g for 5 min.

##### Animals and vascular organoid transplantation

Male and female NOD.Cg-Prkdc^scid^ Il2rg^tm1Wjl^/SzJ (NSG) mice aged 2 to 3 months were obtained from the UCLA Humanized Mouse & Gene Therapy Core.

Each mouse received organoid grafts at two anatomical sites: the ear pinna and dorsal skin. Fifteen day-15 organoids were implanted into the ear pinna, while 50 day-15 organoids were implanted into the dorsal subcutaneous tissue.

Animals were maintained under a 12-h light/12-h dark cycle with unrestricted access to food and water. All animal experiments were approved by the University of California, Los Angeles Chancellor’s Animal Research Committee (IACUC protocol ARC-2025-043).

Ear pinna transplantation. Ear pinna implantation was performed following the method described by Meehan et al. (*57*), with a modification in the position of the surgical incision.

Mice were anesthetized with 2% isoflurane delivered in oxygen at 1.5 L/min. Anesthetic depth was confirmed by the absence of pedal withdrawal responses and monitored through respiratory rate and pattern. Body temperature was maintained using an external heating pad.

The ear pinna was cleaned with 70% ethanol and secured to the barrel of a 5-mL syringe using double-sided adhesive tape. Sterile saline (50 to 100 µL) was injected to create a subcutaneous pocket. An incision was made near the base of the ear pinna adjacent to the head, rather than at the peripheral ear margin.

The opening was enlarged using microdissection scissors, and curved forceps were used to gently separate the skin layers and expand the pocket. Fifteen organoids suspended in Matrigel were placed within the pocket, and the incision was closed using Vetbond tissue adhesive (3M) without sutures.

Dorsal subcutaneous transplantation. Dorsal transplantation was performed in the same recipient mice under inhaled isoflurane anesthesia. The dorsal surgical site was depilated, aseptically prepared, and draped. Lidocaine (0.5%) was administered locally by subcutaneous infiltration before a 0.5 to 1-cm midline dorsal incision was created.

A subcutaneous pocket was generated by blunt dissection using forceps. Fifty organoids suspended in Matrigel were transferred into the pocket using a sterile pipette tip, and the incision was closed with sutures.

Following surgery, animals were transferred to a warmed recovery cage and monitored until fully ambulatory. Mice were assessed twice daily during the first three postoperative days and daily thereafter. Monitoring included inspection of transplantation sites, mobility, grooming behavior, food and water intake, and general appearance.

##### Intravital two-photon microscopy

Intravital two-photon microscopy was performed two months after transplantation to assess vascular organization, local tracer distribution, and immune cell trafficking in ear pinna grafts.

Mice were anesthetized using 2% isoflurane in oxygen delivered at 1.5 L/min. The ear pinna was positioned and immobilized on the imaging platform. PBS was applied to maintain tissue hydration during imaging.

Excitation wavelengths and emission detection channels were selected according to the fluorophores used.

Visualization of endothelial structures. FITC-conjugated Griffonia simplicifolia isolectin B4 (GS-IB4; 20 µg/mL, 5 µL) and ATTO 594-conjugated Ulex europaeus agglutinin I (UEA-I; 2 µg/mL, 5 µL) were administered by local subcutaneous injection near the transplantation site to visualize putative host-derived mouse and graft-associated human endothelial structures, respectively.

Fluorescent dextran tracing. To evaluate local tracer distribution and transport, FITC-conjugated dextran (2,000 kDa; 20 µg/mL, 5 µL) was administered by local subcutaneous injection together with ATTO 594-conjugated UEA-I. Sequential two-photon images were acquired to assess the spatial distribution and movement of the fluorescent tracer relative to UEA-I-labeled structures. Human immune cell trafficking. Human peripheral blood mononuclear cells (PBMCs) were fluorescently labeled. Approximately 5 × 10^5^ labeled PBMCs suspended in 100 µL PBS were administered by local subcutaneous injection at the ear pinna transplantation site.

ATTO 594-conjugated UEA-I was co-administered to visualize graft-associated vascular structures. Intravital two-photon imaging was performed 18 h after PBMC administration to assess immune cell localization, migration, and spatial interactions with UEA-I-labeled structures.

##### Evans blue tracer imaging

Evans blue tracer distribution was evaluated at the ear pinna transplantation site two months after organoid implantation.

Evans blue was prepared at 5 mg/mL in PBS, and 5 µL was administered by local subcutaneous injection at each imaging site.

The distribution of Evans blue was monitored by time-lapse brightfield imaging using an AmScope SM-3B series binocular zoom stereomicroscope (AmScope, Irvine, CA, USA). Images were acquired to document dye movement, distribution, and accumulation at the transplantation site. Illumination, camera exposure, and specimen positioning were maintained consistently across comparable acquisitions.

##### Fluorescent ovalbumin accumulation assay

Organoids were incubated with fluorescent ovalbumin (OVA, Invitrogen) for 12 h. OVA was removed by washing, and organoids were cultured for an additional 3 days before OVA-associated fluorescence was measured.

##### Human fetal tissue procurement

Human fetal tissues were obtained through the UCLA Translational Pathology Core Laboratory (TPCL) following informed consent. Tissue procurement was coordinated through the UCLA Center for Pathology Research Services (CPRS).

##### PBMC incubation with vascular organoids

Human peripheral blood mononuclear cells (PBMCs) were co-cultured with blood vascular organoids (BVOs), lateral plate mesoderm-derived lymphatic organoids (LLOs), and paraxial mesoderm-derived lymphatic organoids (PLOs) to assess their interactions with immune cells. PBMCs were added to the organoid cultures and incubated for 12 h at 37°C. Following incubation, the organoids were gently washed to remove unbound PBMCs and cultured for an additional 12 h. The organoids were then collected, fixed, and subjected to immunofluorescence staining for CCR7. Organoid-associated CCR7-positive cells were visualized by fluorescence microscopy, and CCR7-associated fluorescence intensity was quantified and compared among BVOs, LLOs, and PLOs (Fig. 6, A and B).

#### Histological and molecular characterization

##### Immunofluorescence staining of vascular organoids

Cultured vascular organoids were fixed in 4% paraformaldehyde (PFA) for 30 min at room temperature, washed with phosphate-buffered saline (PBS), and cryoprotected in 30% sucrose overnight at 4°C. Organoids were then embedded in optimal cutting temperature (OCT) compound and cryosectioned at a thickness of 50 µm.

Cryosections were washed with PBS to remove residual OCT and permeabilized and blocked with PBS containing 0.3% Triton X-100 and 5% normal donkey serum for 1 h at room temperature. Sections were incubated with primary antibodies diluted in PBS containing 1% donkey serum and 0.1% Triton X-100 overnight at 4°C. Following three washes with PBS for 10 min each, sections were incubated with species-appropriate fluorophore-conjugated secondary antibodies (1:500) for 2 h at room temperature, protected from light. Sections were subsequently washed three times with PBS, counterstained with DAPI (1 µg/mL) for 5 min, and mounted using an antifade mounting medium. Fluorescence images were acquired using confocal microscopy.

##### Whole-mount immunofluorescence staining

Vascular networks and organoids were fixed in 4% paraformaldehyde (PFA) in Dulbecco’s phosphate-buffered saline (DPBS) for 12 to 14 h at 4°C, washed three times with DPBS, and stored at 4°C until staining. Samples were permeabilized and blocked for 4 h at room temperature with gentle shaking in blocking buffer containing 3% fetal bovine serum (FBS), 1% bovine serum albumin (BSA), 0.5% Triton X-100, 0.5% Tween-20, and 0.01% sodium deoxycholate in PBS. Samples were then incubated with primary antibodies diluted in blocking buffer overnight at 4°C with gentle shaking. Following three washes with DPBS containing 0.05% Tween-20, samples were incubated with fluorophore-conjugated secondary antibodies (1:300) and DAPI (1:500) diluted in blocking buffer for 5 h at room temperature with gentle shaking, protected from light. Samples were subsequently washed three times with DPBS and transferred to 88% glycerol for imaging.

Primary antibodies, suppliers, catalog numbers, and working dilutions are listed in Table S2.

##### Confocal microscopy and image acquisition

Fluorescence images were acquired using Zeiss LSM 880 and LSM 900 confocal laser scanning microscopes.

Laser excitation wavelengths, detector settings, pinhole size, z-step intervals, and image acquisition parameters were selected according to the fluorophores and imaging objectives. Equivalent acquisition settings were maintained across samples included in the same quantitative comparison.

For *in vitro* morphometric analysis, three imaging fields were acquired from each organoid, and three organoids were analyzed per experimental group, resulting in nine fields per group.

##### Endothelial junction morphology quantification

Endothelial junction morphology was quantified from VE-cadherin immunofluorescence images using Fiji/ImageJ.

Images were processed to obtain binary representations of VE-cadherin-positive junctional structures. Segmented images were skeletonized, and individual connected junction objects were identified based on their skeleton identifiers.

For each connected junction object, the lengths of branches sharing the same skeleton identifier were summed to determine its total junction length.

Junctions were operationally classified according to total skeleton length as button-like (≤5 µm), intermediate (>5 to <10 µm), or zipper-like (≥10 µm). Image calibration was verified before length measurements.

The proportional representation of each junction category was calculated as the cumulative junction length assigned to that category divided by the total measured junction length in the image, multiplied by 100.

Mean junction length was calculated from the lengths of individual connected junction objects.

##### Vessel diameter quantification

Vessel diameter was measured using the ImageJ measurement tool based on VE-cadherin-positive endothelial junctions. Measurements represented the external vessel diameter, with six measurements collected per field of view at distinct vessel segments. Measurement positions were selected along clearly defined vessel boundaries, avoiding branch points and regions of overlapping vasculature. For vessels oriented obliquely within the plane, measurements were taken perpendicular to the VE-cadherin-positive vessel boundaries to minimize overestimation of vessel diameter. Measurements from three fields of view were combined within each organoid to calculate an organoid-level mean. Group comparisons were subsequently performed using the organoids rather than individual fields or vessel segments as the sampling units.

##### Transmission electron microscopy

Organoid samples were washed with PBS and fixed overnight at 4 °C in freshly prepared 2% PFA and 2.5% glutaraldehyde in 0.1 M sodium cacodylate buffer (pH 7.4).

Following primary fixation, samples were processed for transmission electron microscopy. Ultrathin sections were collected onto electron microscopy grids and contrasted before imaging.

##### RNA extraction and reverse transcription

Total RNA was extracted from vascular organoids using FreeZol reagent (Vazyme, Cat. No. R711). For each sample, three to four organoids were collected in a 1.5-mL microcentrifuge tube. After removal of the culture medium, 500 µL FreeZol reagent was added. Samples were incubated at room temperature with intermittent vortexing until the organoids were completely lysed. RNA extraction proceeded according to the manufacturer’s protocol.

For reverse transcription, RNA was diluted to 100 ng/µL. Reverse transcription was carried out using the HiScript IV RT SuperMix for qPCR (+gDNA wiper) kit (Vazyme, Cat. No. R423-01) according to the manufacturer’s protocol.

##### Quantitative real-time PCR

Quantitative real-time PCR (RT-qPCR) was performed using SupRealQ Ultra Hunter SYBR qPCR Master Mix (Vazyme, Cat. No. Q713) according to the manufacturer’s protocol. Reactions were run on a real-time PCR detection system (Bio-Rad, Cat. No. 185-5195). Transcript expression was normalized to *GAPDH* using the ΔCq method, where ΔCq = Cq(target) − Cq(*GAPDH*). Normalized relative expression was calculated as 2^−ΔCq^. Primer sequences for *GAPDH*, *TBX6*, *PAX3*, *ETV2*, *CDH5*, *NR2F2*, *PROX1*, *NRP2*, *APLNR*, *EPHB4*, *EFNB2*, *DLL4*, *SOX17*, *PDGFRB*, *FLT4*, *PDPN*, *LYVE1*, *CCL21*, *RELN*, *GATA2*, *ITGA9*, *FOXF1*, and *HAND1* are provided in Table S1.

##### Flow cytometry of the endogenous PROX1 reporter

Organoids were dissociated into single-cell suspensions for flow cytometry. Sequential gates selected cells by forward and side scatter, followed by singlets, live DAPI-negative cells, and CD31-positive endothelial cells. PROX1-mScarlet reporter-positive fractions were calculated within the CD31-positive population (Fig. 2O; Fig. 3O; Fig. S6).

##### Single-cell transcriptomic analysis of vascular organoids

Time-resolved single-cell RNA-sequencing data from BVOs, LLOs, and PLOs were integrated to compare mesodermal and endothelial populations before lymphatic induction. The comparable developmental window contained 110,192 cells (Fig. 4, B to F). Endothelial cells were analyzed separately to compare immature, arterial, angiogenic, and venous states at matched intervals after endothelial induction (Fig. 4, G to I).

Endpoint analyses included 46,284 cells from the three organoid systems. The endothelial compartment comprised 18,864 cells and was reclustered to identify endothelial states (Fig. 5, A to H). Gene-expression and enrichment analyses compared corresponding LLO and PLO lymphatic endothelial states (Fig. 5, I and J).

##### Reference mapping and endothelial classification

Organoid endothelial profiles were compared with human endothelial references spanning tissue types and developmental stages. Reference-based label transfer was used to assign endothelial identities. Discriminant-space projections compared organoid states with tissue and developmental-stage reference groups (Fig. 5, K to N). Seurat canonical-correlation-based classification distinguished LLO and PLO lymphatic endothelial cells. Held-out organoid cells and human embryonic lymphatic endothelial cells were evaluated using LLO-class prediction scores (Fig. 5O).

##### Predicted endothelial-stromal communication

CellChat (49) was used to infer ligand-receptor interactions between supporting-cell populations and endothelial states in LLOs and PLOs. Analyses compared predicted communication networks, ligand-receptor pairs, and relative ligand-associated contributions from stromal populations (Fig. S8, G to K). Exported CellChat tables were used to generate the signaling panels.

##### Statistical analysis

All statistical tests were two-sided unless otherwise specified.

For spatial transcriptomic analyses, adjacency fractions were compared using Fisher’s exact tests. Pseudotime distributions were compared using Mann-Whitney tests. No multiple-testing correction was applied to the two prespecified spatial comparisons, and the underlying cell counts and proportions were reported with the corresponding Figures.

For *in vitro* organoid measurements, data from individual imaging fields and junction objects were aggregated at the organoid level before statistical testing. Individual fields and junction objects were considered subsamples and not independent biological replicates.

##### Software

Single-cell transcriptomic analyses were performed using Python 3.11.15 with anndata 0.12.19, NumPy 2.4.6, pandas 2.3.3, SciPy 1.16.3, scikit-learn 1.9.0, harmonypy 2.0.0, and Matplotlib 3.11.0.

R-based analyses used Seurat 5.4.0, SingleCellExperiment 1.30.1, Monocle 2.36.0, Slingshot 2.18.0, and Harmony 1.2.4.

The reported package versions correspond to the analysis environment used to produce the Figures.

Image processing and morphometric analyses were performed using Fiji/ImageJ.

---

**Fig. S1.**
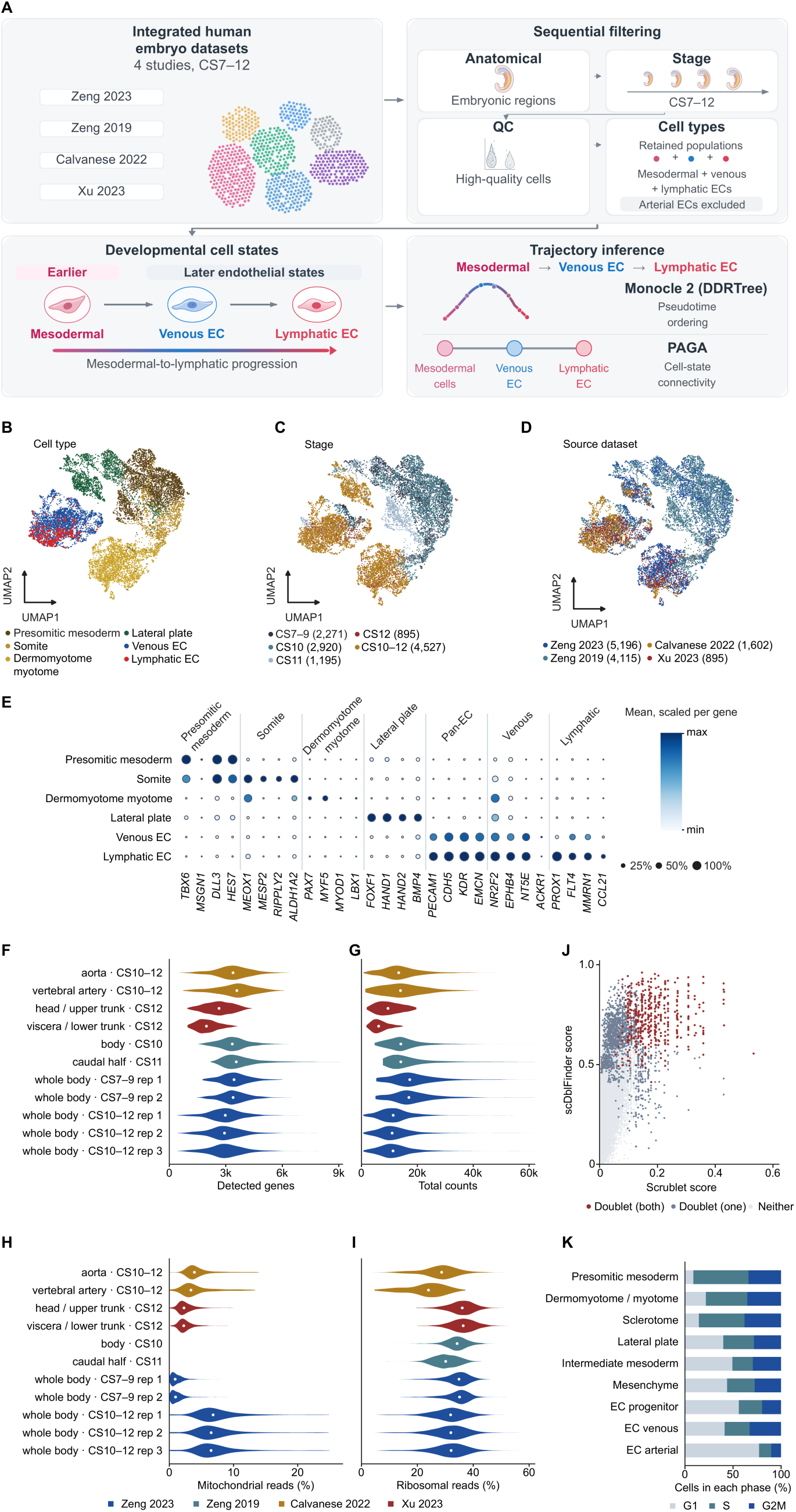
Embryonic reference integration, cell annotation, and quality control. **(A)** Schematic of assembly of the human embryonic single-cell reference from four studies covering CS7 to CS12, sequential filtering for relevant embryonic anatomical regions and mesodermal/venous/lymphatic cell types, followed by quality control and trajectory inference. Arterial endothelial populations were not included in the final mesoderm-to-lymphatic trajectory models. **(B)** UMAP of integrated human embryonic mesodermal and endothelial cells colored by the six displayed cell-type classes. **(C)** The same UMAP colored by source developmental stage or stage interval (CS7 to 9, CS10, CS11, CS12, and CS10 to 12). Cells assigned only to a stage interval were retained as such, rather than forced into a single Carnegie stage. **(D)** The same UMAP colored by contributing study: Zeng et al. (*24*) (n = 5,196); Zeng et al. (*27*) (n = 4,115); Calvanese et al. (*28*) (n = 1,602); and Xu et al. (*29*) (n = 895). **(E)** Dot plot of representative identity markers across presomitic mesoderm, somite, dermomyotome, lateral plate mesoderm, venous endothelium, and lymphatic endothelium. Dot size represents the percentage of marker-detecting cells and color denotes gene-wise scaled mean expression. **(F to I)** Quality-control distributions by contributing dissection or sample: **(F)** number of detected genes, **(G)** total transcript/UMI counts, **(H)** mitochondrial read fraction, and **(I)** ribosomal read fraction. Violin plots show the distributions for each contributing sample. **(J)** Joint display of Scrublet and scDblFinder doublet scores, colored by whether cells were called doublets by both methods, one method, or neither method. **(K)** Proportions of cells assigned to G1, S, or G2/M phases within the indicated embryonic cell populations. Values are percentages of each population.

**Fig. S2.**
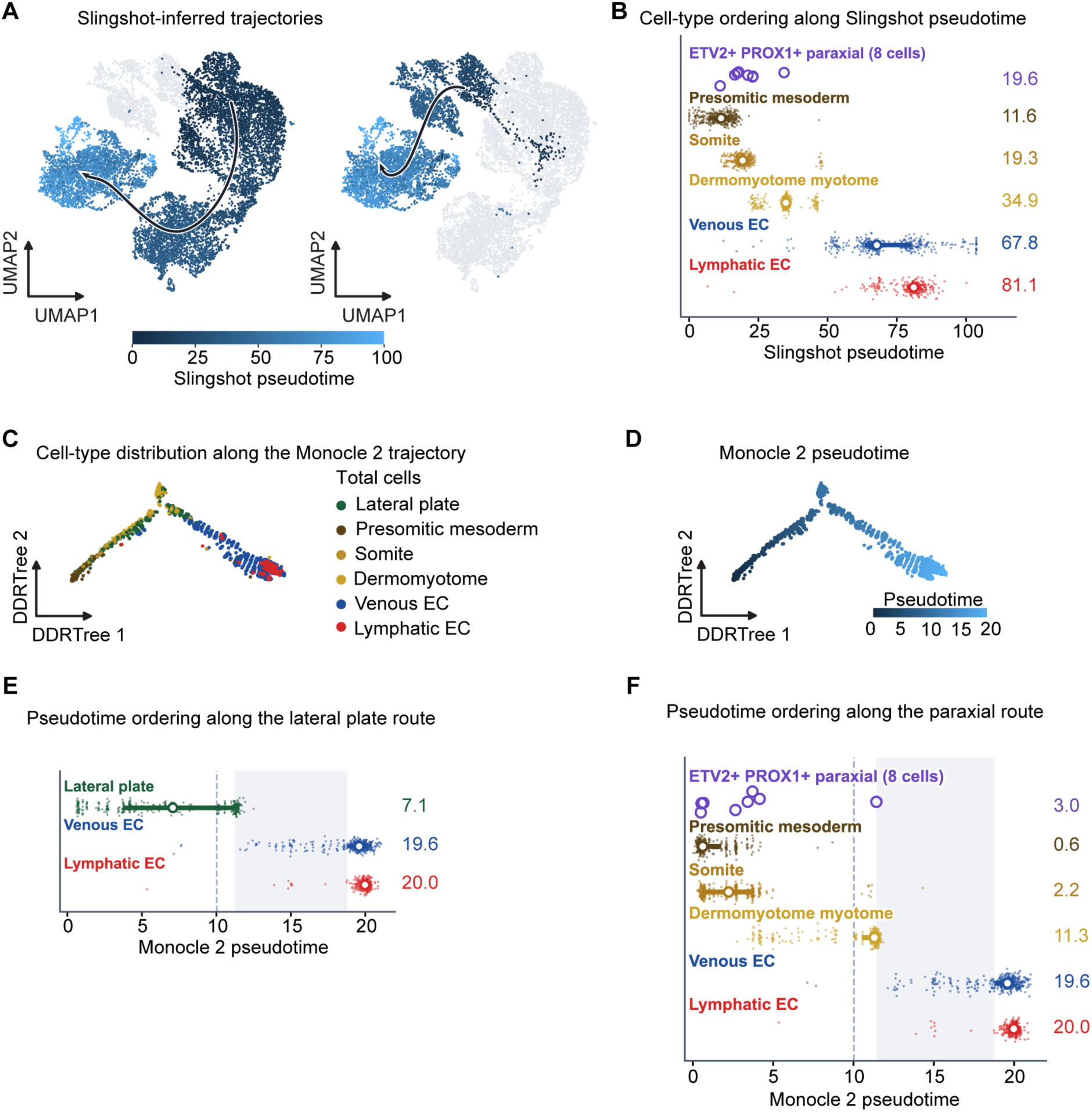
Independent pseudotime reconstructions and the position of ETV2⁺PROX1⁺ paraxial cells. **(A)** Slingshot analysis of the paraxial route on the integrated embryonic UMAP. The inferred principal trajectory is shown on the selected route cells, together with a view colored by route-specific Slingshot pseudotime (rescaled from 0 to 100). **(B)** Pseudotime ranges for presomitic mesoderm, somite, dermomyotome/myotome, venous endothelial cells, and lymphatic endothelial cells; horizontal bars span the 10th to 90th percentile within each class. Purple symbols mark the positions of ETV2⁺PROX1⁺ paraxial cells. **(C)** Monocle 2 DDRTree projection of the route-colored cell populations. **(D)** The same DDRTree embedding colored by Monocle 2 pseudotime. **(E)** Distribution of Monocle 2 pseudotime for lateral plate mesoderm, venous endothelial cells, and lymphatic endothelial cells. Points represent cells and open circles indicate the displayed median pseudotimes; the vertical dashed line and shaded band mark the indicated mesoderm-to-endothelial transition range. **(F)** Monocle 2 pseudotime distributions for paraxial-route cell populations, displayed as in (E), with ETV2⁺PROX1⁺ paraxial cells shown as a separate category.

**Fig. S3.**
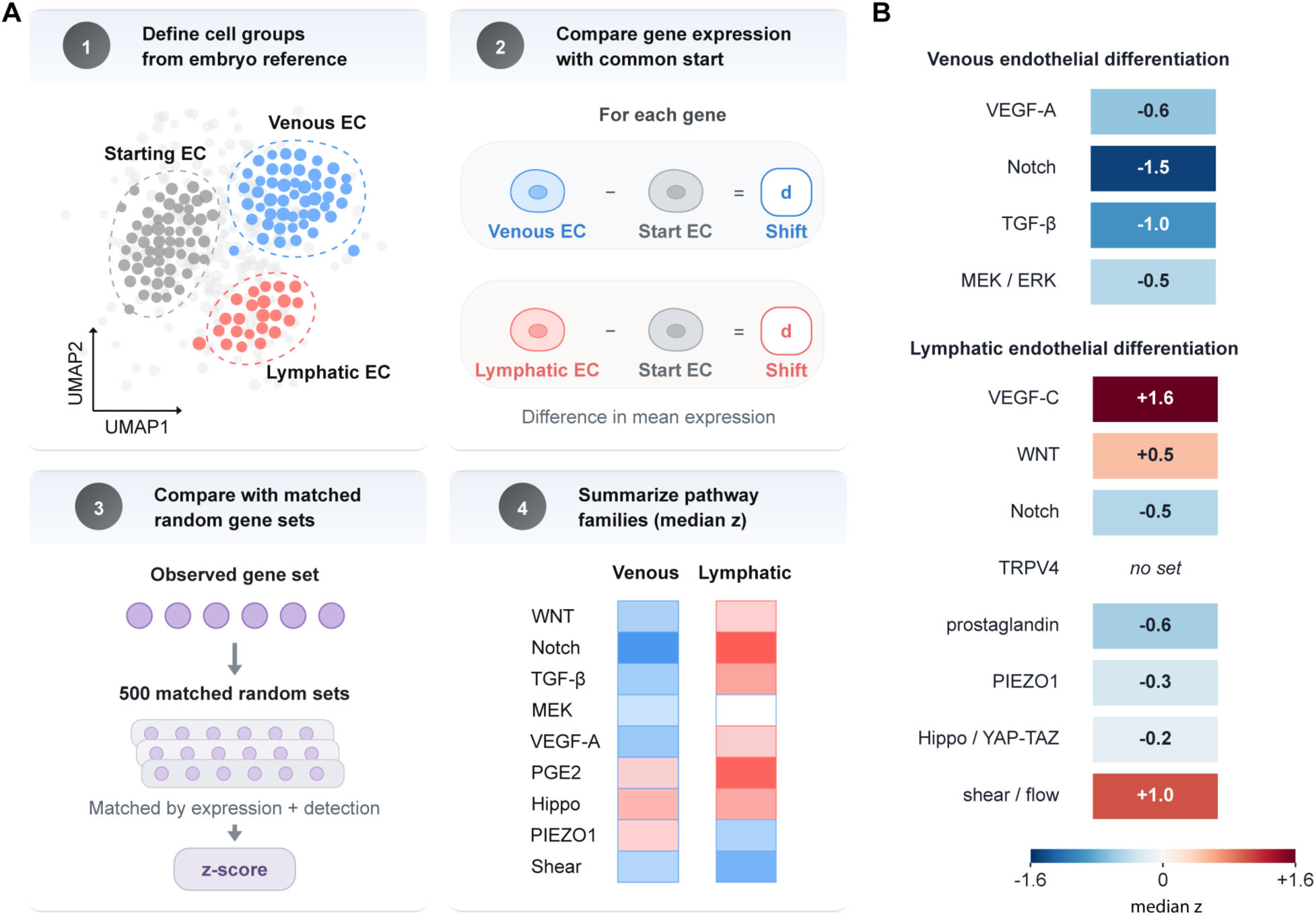
Transcriptome-based pathway prioritization for venous and lymphatic endothelial induction. **(A)** Overview of the pathway-prioritization analysis. Starting endothelial/progenitor, venous endothelial, and lymphatic endothelial populations were selected from the embryonic reference. For each gene, expression differences from the common starting population were calculated independently for the venous and lymphatic comparisons. Gene-set shifts were evaluated using 500 random gene sets matched for expression level and detection frequency, converted to standardized scores, and summarized by pathway family using the median score. The analysis used the full curated MSigDB 2023.2.Hs collection of 8,903 gene sets; it did not treat individual curated sets as independent pathway activity measurements. **(B)** Signed pathway-family scores associated with venous specification (VEGFA, −0.6; NOTCH, −1.5; TGF-β, −1.0; and MEK/ERK, −0.5) and lymphatic specification (*VEGFC*-related, +1.6; WNT, +0.5; NOTCH, −0.5; prostaglandin, −0.6; PIEZO1, −0.3; Hippo/YAP-TAZ, −0.2; and shear/flow, +1.0). “No set” indicates that no qualifying curated gene set was available. Scores reflect expression-associated gene-set patterns standardized against matched random sets.

**Fig. S4.**
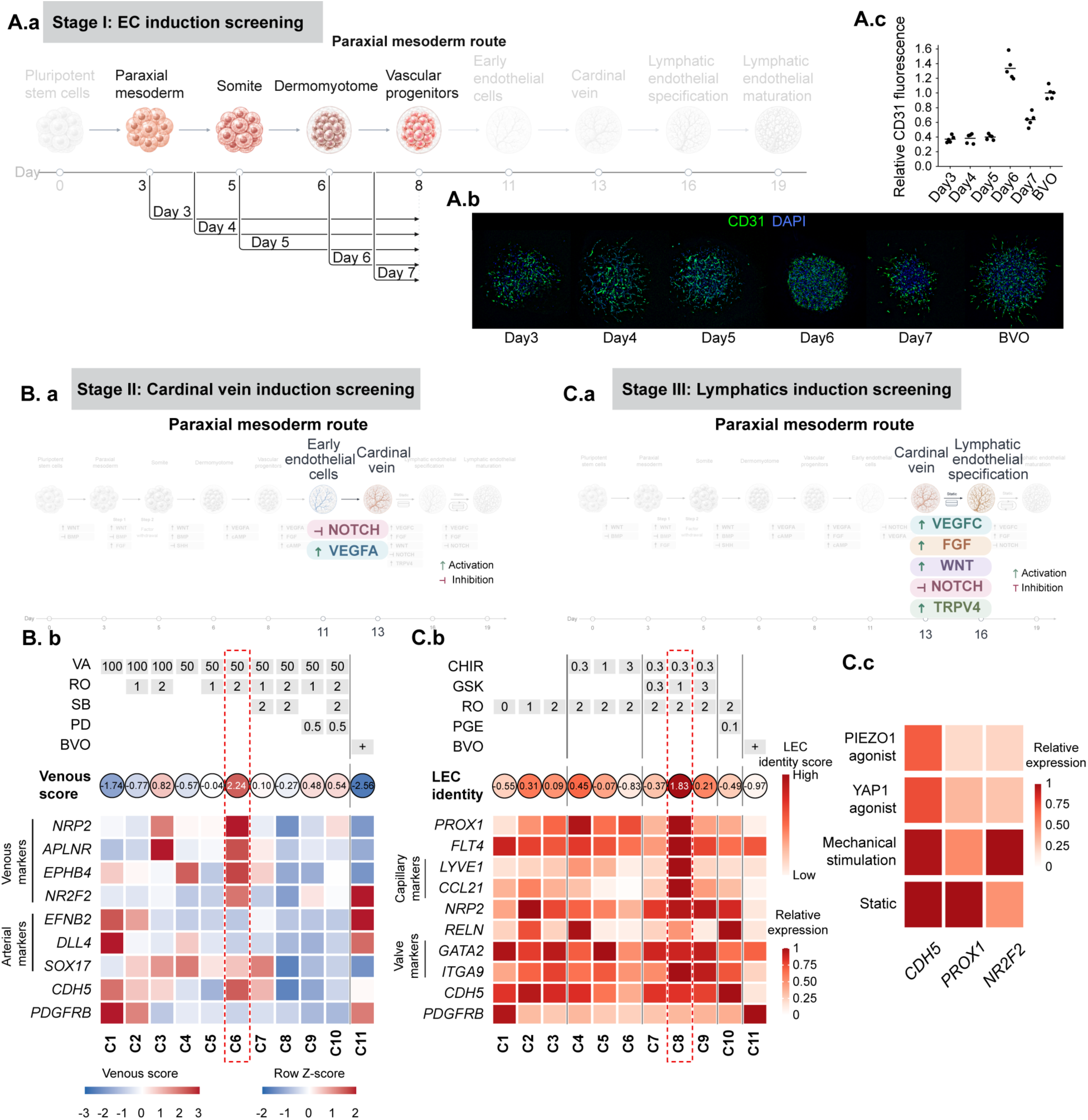
Stepwise optimization of paraxial endothelial, cardinal vein-like, and lymphatic induction. (A.a) Stage I endothelial induction screening along the paraxial mesoderm route, showing endothelial induction initiated on successive days 3 to 7. **(A.b)** Representative immunofluorescence images of CD31 (green) and DAPI (blue) under the indicated induction conditions, with BVOs as a reference. **(A.c)** Quantification of relative CD31 fluorescence intensity under the indicated conditions, showing the highest signal following endothelial induction initiated on day 6. **(B.a)** Stage II cardinal vein-like endothelial induction screening, showing the intervention window from early endothelial cells to a cardinal vein-like endothelial state. **(B.b)** Comparative venous induction screen using different concentrations of VEGFA (VA) and combinations with NOTCH inhibition (RO), TGF-β pathway inhibition (SB), and MEK/ERK pathway inhibition (PD), with BVOs as a reference (C11). Circles indicate composite venous scores, calculated as the difference between the median scores of venous-associated markers (*NRP2*, *APLNR*, *EPHB4*, *NR2F2*) and arterial markers (*EFNB2*, *DLL4*, *SOX17*). Heatmap colors represent per-gene row z-scores, with *CDH5* and *PDGFRB* included as endothelial and mural reference markers, respectively. Condition C6 yielded the highest venous score (2.24) among the tested conditions. **(C.a)** Stage III lymphatic endothelial induction screening following the cardinal vein-like stage, showing the intervention window for lymphatic endothelial specification. **(C.b)** Comparative lymphatic induction screen using combinations of WNT activation (CHIR), TRPV4 activation (GSK), NOTCH inhibition (RO), and prostaglandin treatment (PGE), with BVOs as a reference (C11). Circles indicate lymphatic endothelial identity scores, and heatmap colors represent relative gene expression. The heatmap includes lymphatic and capillary-associated markers (*PROX1*, *FLT4*, *LYVE1*, *CCL21*, *NRP2*, *RELN*), valve-associated markers (*GATA2*, *ITGA9*), and endothelial/mural reference markers (*CDH5*, *PDGFRB*). Condition C8, combining WNT activation, TRPV4 activation, and NOTCH inhibition, yielded the highest lymphatic endothelial identity score (1.83) among the tested conditions. **(C.c)** Heatmap comparing relative expression of *CDH5*, *PROX1*, and *NR2F2* under static culture, PIEZO1 agonism, YAP1 agonism, and mechanical stimulation. Heatmap colors indicate relative gene expression levels.

**Fig. S5.**
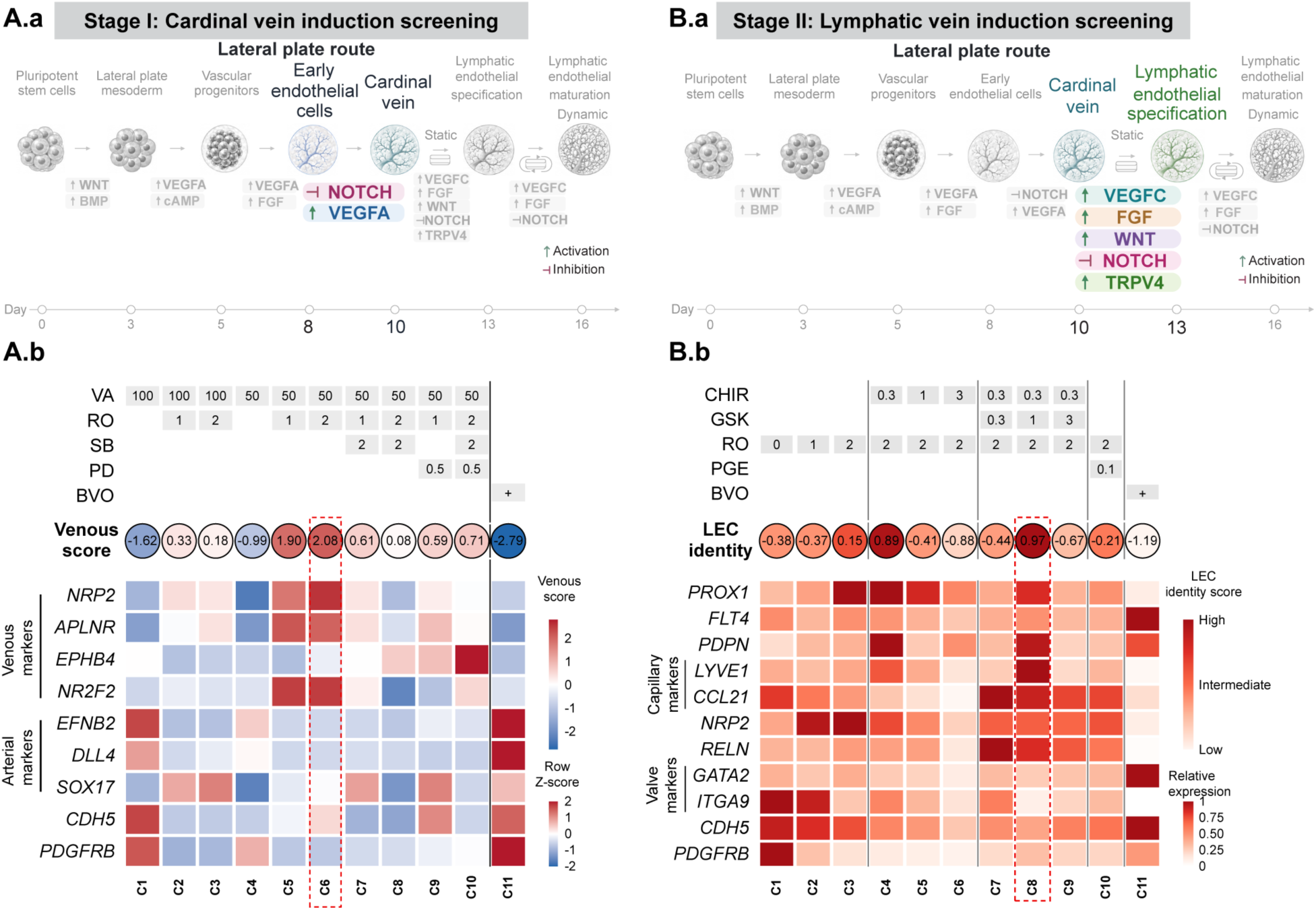
Venous and lymphatic induction screens in the lateral plate mesoderm differentiation route. (**A.a**) Stage I cardinal vein-like endothelial induction screening along the lateral plate mesoderm route, showing the intervention window from early endothelial cells to a cardinal vein-like endothelial state. (**A.b**) Comparative venous induction screen using different concentrations of VEGFA (VA) and combinations with NOTCH inhibition (RO), TGF-β pathway inhibition (SB), and MEK/ERK pathway inhibition (PD), with BVOs as a reference (C11). Circles indicate composite venous scores, calculated as the difference between the median expression scores of venous-associated markers (NRP2, APLNR, EPHB4, NR2F2) and arterial markers (EFNB2, DLL4, SOX17). Heatmap colors represent per-gene row z-scores, with CDH5 and PDGFRB included as endothelial and mural reference markers, respectively. Condition C6 yielded the highest venous score (2.08) among the tested lateral plate-route conditions. (**B.a**) Stage II lymphatic endothelial induction screening following the cardinal vein-like stage, showing the intervention window for lymphatic endothelial specification. (**B.b**) Comparative lymphatic induction screen using combinations of WNT activation (CHIR), TRPV4 activation (GSK), NOTCH inhibition (RO), and prostaglandin treatment (PGE), with BVOs as a reference (C11). Circles indicate lymphatic endothelial identity scores, and heatmap colors represent relative gene expression. The heatmap includes lymphatic and capillary-associated markers (PROX1, FLT4, PDPN, LYVE1, CCL21, NRP2, RELN), valve-associated markers (GATA2, ITGA9), and endothelial/mural reference markers (CDH5, PDGFRB).

**Fig. S6.**
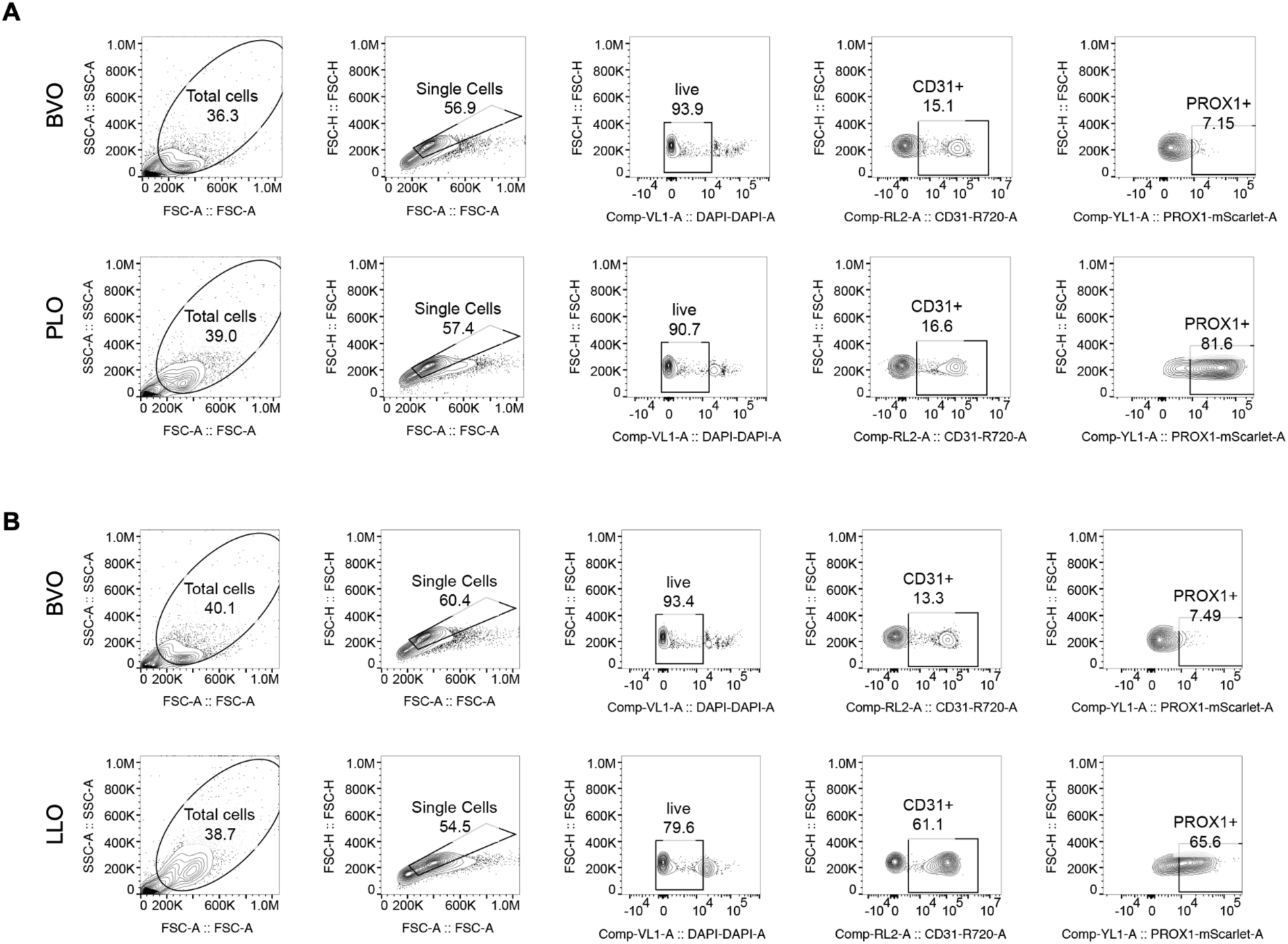
Representative flow-cytometric gating of endogenous PROX1 reporter expression. **(A)** Sequential flow-cytometry gating for BVO and PLO samples: total events by forward/side scatter, singlet discrimination, live DAPI-negative cells, CD31-positive endothelial cells, and PROX1-mScarlet reporter-positive cells. The final percentages of PROX1 reporter-positive cells among CD31⁺ cells are 7.15% in BVOs and 81.60% in PLOs, corresponding to Fig. 2O. **(B)** Equivalent gating workflow for BVO and LLO samples. The final proportions of PROX1 reporter-positive cells among CD31⁺ cells are 7.49% in BVOs and 65.60% in LLOs, corresponding to Fig. 3O. Percentages shown at earlier gating steps are conditional on each respective parent gate.

**Fig. S7.**
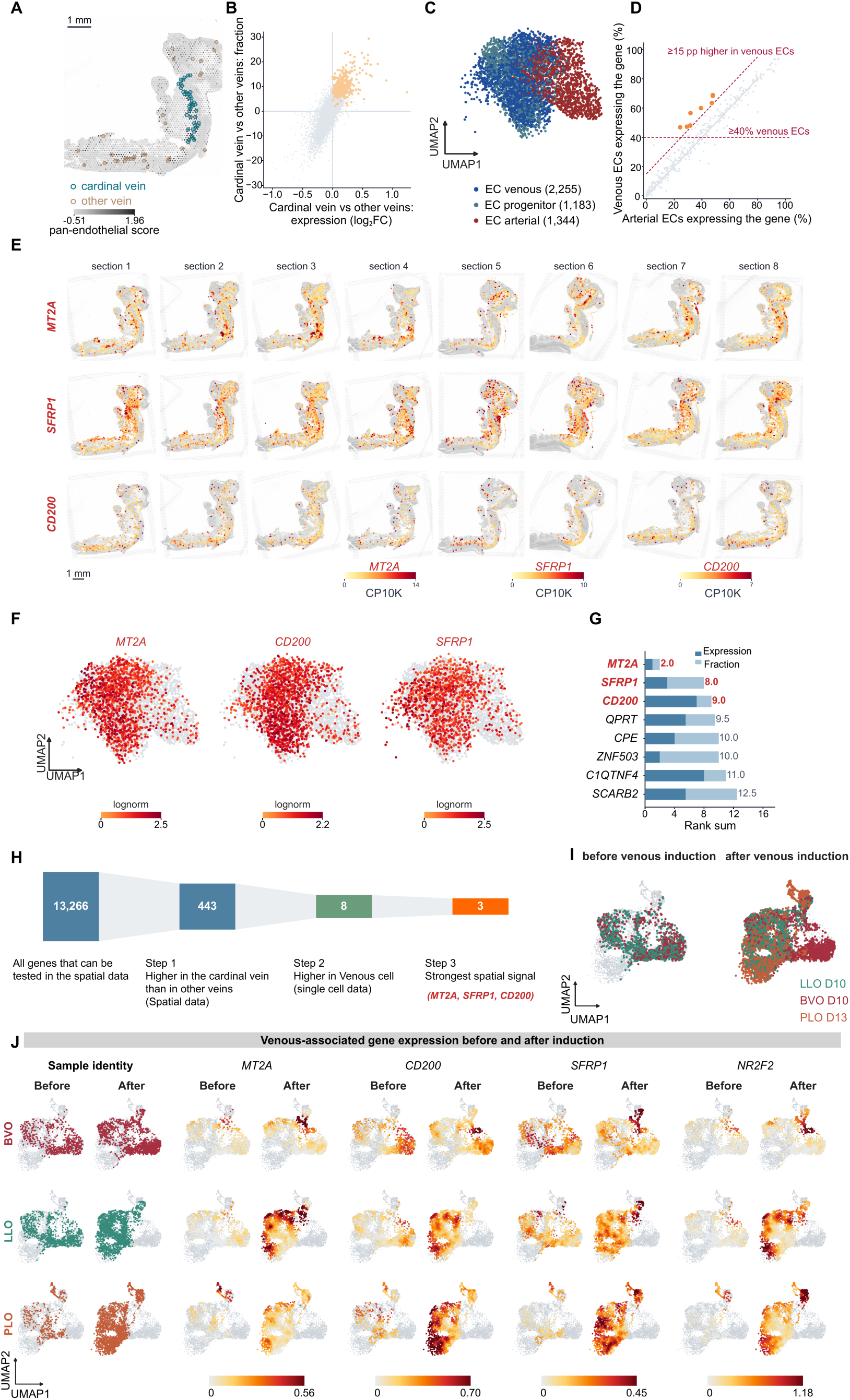
Integrated spatial and single-cell screening identifies candidate cardinal vein-associated markers. **(A)** Manually annotated candidate cardinal-vein and other venous regions in a CS12 human embryo spatial section. Open cyan circles: candidate cardinal-vein capture spots; brown circles: spots from other venous territories. Anatomical position and endothelial/venous-marker expression informed the vein annotations. Scale bar, 1 mm. **(B)** Spatial comparison of candidate cardinal-vein versus other-vein expression for genes testable in the spatial dataset. The horizontal axis shows expression fold change (log2), and the vertical axis shows the difference in detection fraction; orange points: genes retained at first selection step. **(C)** Endothelial-only UMAP from the corresponding embryonic single-cell reference, with venous endothelial (n = 2,255), endothelial progenitor (n = 1,183), and arterial endothelial (n = 1,344) populations. **(D)** Comparison of detection frequencies in venous versus arterial endothelial cells for the spatially enriched candidates. Eight selected genes met the indicated criteria: detection in at least 40% of venous cells and ≥15 percentage points greater detection than in arterial cells. **(E)** Spatial expression maps for *MT2A*, *SFRP1*, and *CD200* across all eight embryonic tissue sections, showing gene expression at the corresponding spatial capture spots. Color scales are gene-specific and expressed as counts per 10,000 transcripts (CP10K). **(F)** Expression of *MT2A*, *CD200*, and *SFRP1* in the endothelial UMAP from (C). Colors represent log-normalized gene expression, scaled separately for each marker. **(G)** Relative ranking of eight shortlisted genes based on spatial enrichment and expression-based criteria. *MT2A*, *SFRP1*, and *CD200* were selected as the three final candidate markers. **(H)** Summary of the screening process: 13,266 genes testable in the spatial data, 443 spatially enriched candidates, eight genes with preferential venous endothelial detection, and three markers prioritized by spatial expression (*MT2A*, *SFRP1*, *CD200*). **(I)** Integrated endothelial UMAP displaying before- and after-venous-induction samples from BVOs (days 8 and 10), LLOs (days 8 and 10), and PLOs (days 11 and 13), distinguishing sample identity by color. **(J)** Gene-expression feature maps for *MT2A*, *CD200*, *SFRP1*, and *NR2F2* in the same BVO, LLO, and PLO samples before and after venous induction. Color indicates expression relative to the gene-specific scale; the maps are descriptive and do not imply equivalent expression ranges across different genes.

**Fig. S8.**
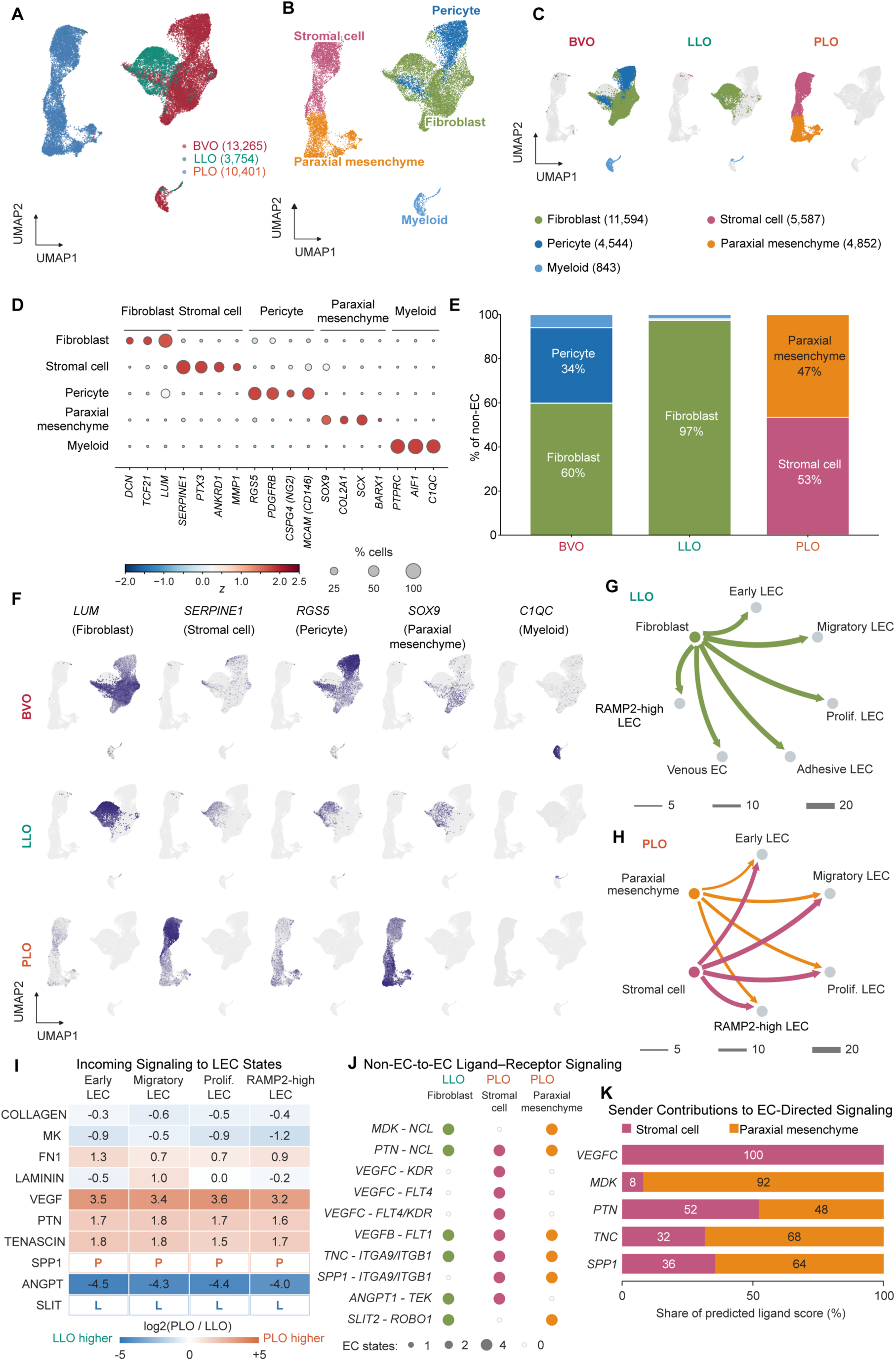
Route-specific stromal populations and predicted endothelial-stromal signaling in lymphatic organoids. **(A)** Integrated UMAP of non-endothelial cells from BVOs (n = 13,265), LLOs (n = 3,754), and PLOs (n = 10,401), colored by organoid source (27,420 cells total). **(B)** The same non-endothelial UMAP colored by fibroblast (n = 11,594), stromal cell (n = 5,587), pericyte (n = 4,544), paraxial mesenchyme (n = 4,852), and myeloid (n = 843) populations. **(C)** The same embedding split by BVO, LLO, and PLO groups, with cells colored by cell class and the remaining cells shown in gray. Cell numbers above the panels indicate cells assigned to each organoid. **(D)** Dot plot of genes identifying the annotated non-endothelial populations, including fibroblast-associated extracellular matrix genes, stromal markers, pericyte-associated genes, paraxial mesenchymal markers, and myeloid genes. Dot size is the percentage of cells expressing the indicated gene and fill represents per-gene scaled mean expression. **(E)** Non-endothelial composition by organoid type. The dominant displayed classes were fibroblasts and pericytes in BVOs (60% and 34%), fibroblasts in LLOs (97%), and stromal cells and paraxial mesenchyme in PLOs (53% and 47%). Values are rounded percentages of the non-endothelial compartment. **(F)** Feature plots for representative non-endothelial markers: *LUM* (fibroblast), *SERPINF1* (stromal cell), *RGS5* (pericyte), *SOX9* (paraxial mesenchyme), and *C1QC* (myeloid), displayed separately for BVOs, LLOs, and PLOs. **(G)** Inferred ligand-receptor communication network from LLO fibroblasts to the indicated endothelial states, including early, migratory, proliferating, RAMP2-high, venous, and pre-collecting-like endothelial categories. Line width denotes the inferred strength/number of ligand-receptor interactions as represented in the Figure. **(H)** Analogous inferred signaling networks from PLO paraxial mesenchyme and stromal cells to lymphatic endothelial states. Edges represent CellChat-predicted ligand-receptor interactions. **(I)** Expression contrasts (log2[PLO/LLO]) for selected extracellular matrix, signaling, and ligand-related gene families in matched lymphatic endothelial states. Positive values denote higher expression in PLO-derived cells and negative values higher expression in LLO-derived cells; letters mark lineage-restricted categories shown in the heatmap. **(J)** Predicted ligand-receptor pairs connecting LLO fibroblasts or PLO stromal/paraxial mesenchymal populations with endothelial states. The pairs include MDK-NCL, PTN-NCL, VEGFC-KDR, VEGFC-FLT4, VEGFC-FLT4/KDR, VEGFB-FLT1, TNC-ITGA9/ITGB1, SPP1-ITGA9/ITGB1, ANGPT1-TEK, and SLIT2-ROBO1. Dot size represents the number of endothelial states supporting each predicted interaction (0, 1, 2, or 4 as shown). **(K)** Estimated contributions of PLO stromal cells and paraxial mesenchyme to the indicated ligand-associated scores. The PLO stromal population accounted for 100% of the displayed VEGFC score, whereas the estimated contributions of the two populations differed for MDK, PTN, TNC, and SPP1.

**Table S1.**
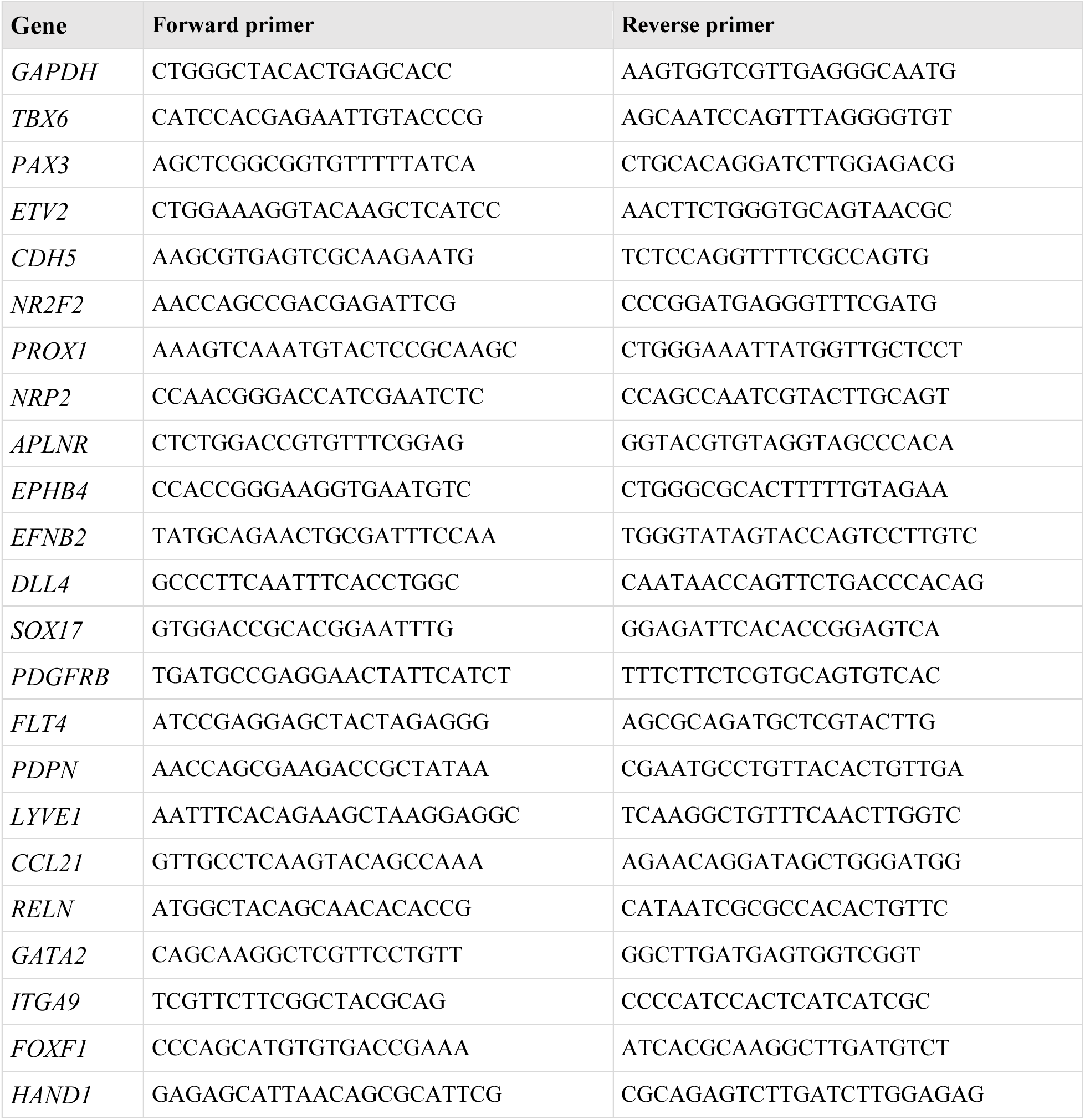
RT-qPCR primer sequences.

**Table S2.** Primary antibodies used for immunofluorescence staining.

| Target | Supplier | Catalog number | Dilution |
|---|---|---|---|
| HAND1 | R&D Systems | AF3168-SP | 1:300 |
| PAX3 | Developmental Studies Hybridoma Bank | Pax3-c | 1:100 |
| VE-cadherin (CDH5) | R&D Systems | AF938 | 1:300 |
| NR2F2 (COUP-TFII) | Abcam | ab41859 | 1:300 |
| PROX1 | Abcam | ab199359 | 1:100 |
| CD31 (PECAM1) | R&D Systems | AF806 | 1:300 |
| PDGFRβ | Abcam | ab32570 | 1:200 |
| LYVE1 | Abcam | ab36993 | 1:300 |
| ITGA9 | Abcam | ab27947 | 1:300 |
| PDPN (podoplanin) | Abcam | ab10288 | 1:300 |
| CCR7 | Abcam | ab253187 | 1:300 |
| CD31-VioBright R720 | Miltenyi Biotec | 130-128-241 | 2ul / test |

**Table S3.**
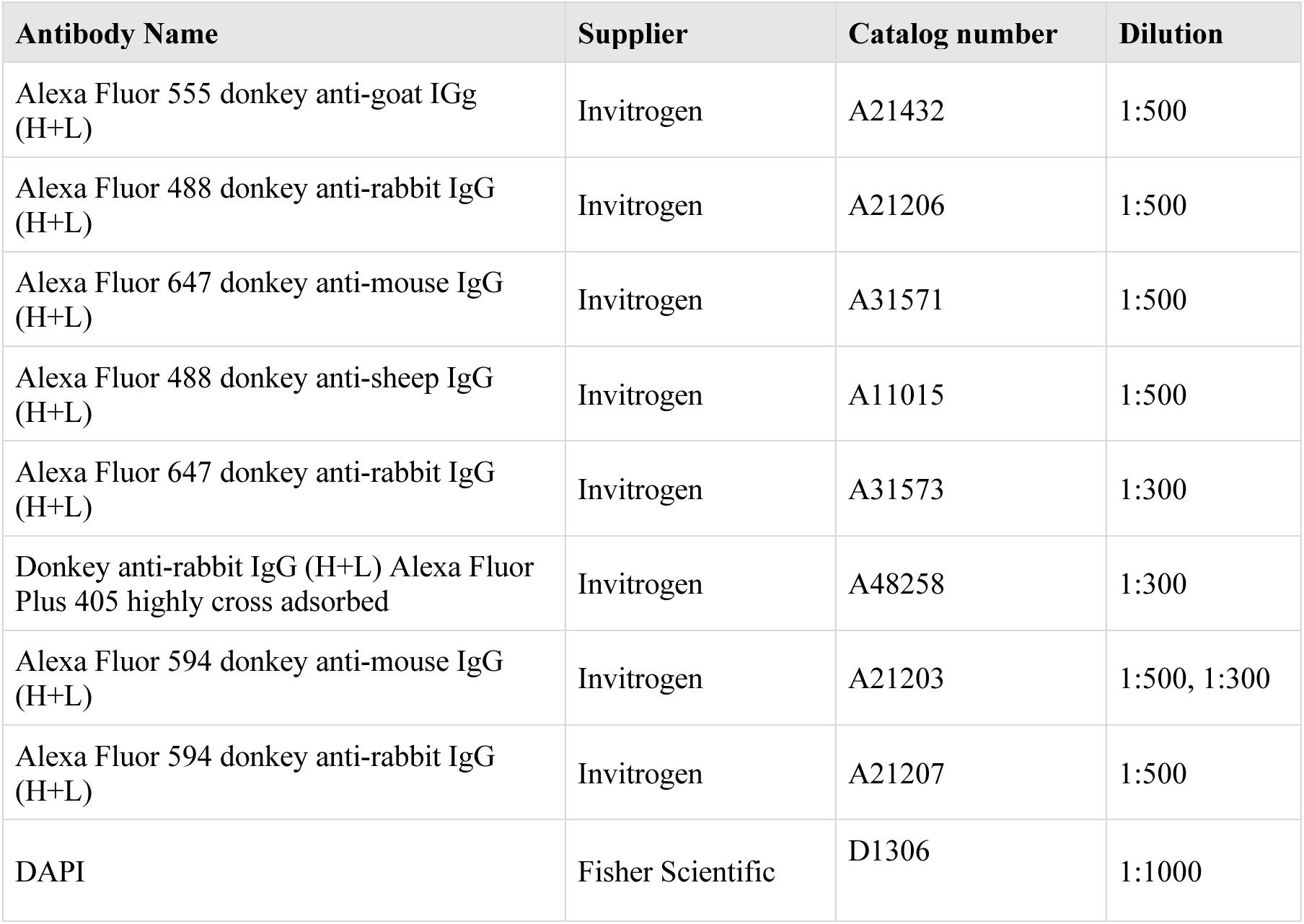
Secondary antibodies and dilutions.

| Antibody Name | Supplier | Catalog number | Dilution |
|---|---|---|---|
| Alexa Fluor 555 donkey anti-goat IGg (H+L) | Invitrogen | A21432 | 1:500 |
| Alexa Fluor 488 donkey anti-rabbit IgG (H+L) | Invitrogen | A21206 | 1:500 |
| Alexa Fluor 647 donkey anti-mouse IgG (H+L) | Invitrogen | A31571 | 1:500 |
| Alexa Fluor 488 donkey anti-sheep IgG (H+L) | Invitrogen | A11015 | 1:500 |
| Alexa Fluor 647 donkey anti-rabbit IgG (H+L) | Invitrogen | A31573 | 1:300 |
| Donkey anti-rabbit IgG (H+L) Alexa Fluor Plus 405 highly cross adsorbed | Invitrogen | A48258 | 1:300 |
| Alexa Fluor 594 donkey anti-mouse IgG (H+L) | Invitrogen | A21203 | 1:500, 1:300 |
| Alexa Fluor 594 donkey anti-rabbit IgG (H+L) | Invitrogen | A21207 | 1:500 |
| DAPI | Fisher Scientific | D1306 | 1:1000 |

